# *TP53*-mutant AML with ribosomal gene loss exhibits impaired protein translation and sensitivity to HSP90 inhibition

**DOI:** 10.1101/2025.10.01.679539

**Authors:** Jean-François Spinella, Jalila Chagraoui, Isabel Boivin, Guillaume Richard Carpentier, Céline Moison, Nadine Mayotte, François Béliveau, Vincent P. Lavallée, Josée Hébert, Guy Sauvageau

## Abstract

*TP53*-mutated acute myeloid leukemia (AML) represents a particularly aggressive and therapeutically refractory subtype of the disease. While recurrent chromosomal abnormalities such as -5/del(5q), -7/del(7q), and del(17p) are well studied in this context, additional co-occurring events remain less well defined. Using the multi-dimensional Leucegene dataset (∼700 primary AML specimens), we identified and comprehensively characterized a distinct subset of *TP53*-altered AML marked by recurrent deletions on the short arm of chromosome 3 (del(3p), > 20% *TP53*-mutated cases). These deletions frequently co-occur with del(5q) and encompass several ribosomal protein genes (RPGs), leading to a global downregulation of the ribosomal network and reduced protein synthesis. We show that this ribosomopathy-like phenotype is most pronounced in *TP53*-mutated cases with combined RPG deletions on chromosomes 3p and 5q, suggesting a cooperative oncogenic mechanism. Importantly, chemical screening identified HSP90 inhibition as a selective vulnerability in AML with low RPG expression. These findings highlight a previously unappreciated *TP53*-altered AML subset characterized by converging genomic and translational defects, and suggest that ribosomal stress may serve as a therapeutic entry point for targeted intervention of this patient subgroup.

## Introduction

Acute myeloid leukemia (AML) is a genetically heterogeneous malignancy characterized by recurrent chromosomal aberrations and gene mutations. Its diagnosis is based on complementary laboratory tests including morphology, flow cytometry, cytogenetics and targeted mutation analysis^1^ that enable classification of this disease into distinct biological entities and allow risk stratification.

Despite regular advances, the clinical outcome of AML remains poor, with a 5-year survival of 32.9%^2^. Despite the development of newer therapeutic agents, long-lasting remissions remain uncommon for high-risk disease^1,3^. Among the most aggressive subtypes is *TP53*-altered AML, which is found in approximately 5-10% of *de novo* cases and up to 30% of therapy-related or elderly AML^4,5^.

AML with mutated *TP53* was classified as a distinct diagnostic entity in the 2022 International Consensus Classification (ICC). Under this definition, any AML case with ≥ 20% blasts and a pathogenic *TP53* mutation exhibiting a variant allele frequency (VAF) of ≥10% qualifies for inclusion in this category^6^. *TP53*-mutated AML shows a unique biology with extremely poor prognosis with high relapse rate post hematopoietic stem cell transplantation^7,8^.

Although *TP53*-mutated AML frequently harbors large chromosomal deletions – most notably of 5q, 7q, and 17p^4,9^ – the landscape of other, potentially cooperating genomic lesions is poorly defined. Given the central role of TP53 in DNA repair, apoptosis, and cellular stress responses^4^, delineating the genomic context in which TP53 loss-of-function drives leukemogenesis is crucial to the identification of novel disease mechanisms and vulnerabilities.

In this study, we used the Leucegene dataset (https://leucegene.ca/), a comprehensive, multi-dimensional resource comprising 691 AML specimens from 656 patients selected to capture the full genetic diversity of the disease. Through integrative transcriptomic and genomic analyses, we identified a subset of *TP53*-altered AMLs characterized by recurrent deletions on chromosome 3p that co-occur with del(5q). These deletions span several ribosomal protein genes (RPGs), resulting in network-wide downregulation of the ribosome and impaired protein synthesis, a phenotype evoking somatic ribosomopathies. This ribosomal dysfunction defines a biologically distinct subgroup of *TP53*-mutated AML, for which we demonstrate increased sensitivity to HSP90 inhibition, highlighting a potential therapeutic vulnerability.

## Results

Accompanying this study, we report the complete Leucegene data collection [https://leucegene.ca/] including genomic and transcriptomic data, as well as clinical annotations (see “**Data Availability**” section), for 691 primary AML specimens. These samples are distributed into 15 cytogenetically defined molecular subgroups, with frequencies ranging from 0.3% (−17/del(17p) not complex and hyperdiploid with numerical abnormalities only) to 40.8% (normal karyotype) (**Figure 1A, left panel, and Sup. Table S1**; see **Figure 1A, right panel**, for the distribution of *TP53*-mutated cases among molecular subgroups; see **Figure 1B** for the distribution of general clinical characteristics).

**Figure 1.**
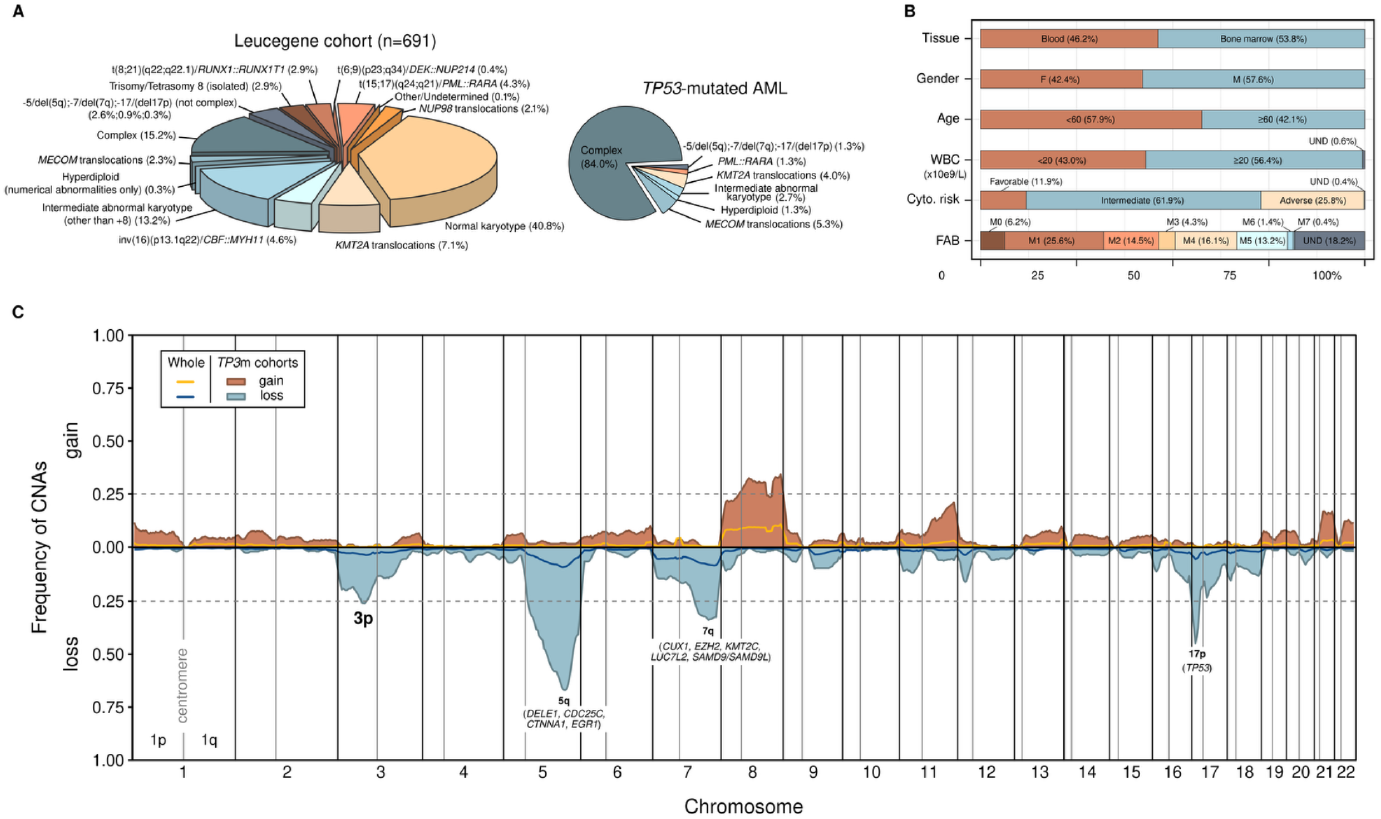
**A**. Frequency of cytogenetic subgroups in the Leucegene cohort (n = 691, left panel) and distribution of *TP53*-mutated cases in these subgroups (right panel). **B**. Distribution of general clinical characteristics of AML specimens in the Leucegene cohort. FAB, French-American-British classification; WBC, white blood cell count; UND, undetermined. **C**. CNA landscape of the whole Leucegene cohort (yellow and dark blue curves for gains and losses, respectively) or limited to *TP53*-mutated AML (brown and light blue shaded area for gains and losses, respectively). Reported mean frequencies of gain (top panel) and loss events (bottom panel) are calculated per window of 3 Mb. Main haploinsufficient candidate genes are indicated under the peaks formed by well-characterized *TP53*m-associated deletions (5q, 7q and 17p). Vertical solid black and gray lines separate chromosomes and depict centromeres, respectively. Horizontal dotted lines indicate 25% of frequency.

We analyzed non-redundant specimens of the collection (n = 656), prioritizing primary diagnostic samples. All specimens were subjected to RNA sequencing (RNA-seq, **Methods**), enabling comprehensive mutational (**Sup. Figure S1**) and transcriptomic profiling. In parallel, we used low-pass whole-genome sequencing (WGS, **Methods**) to assess copy number alterations (CNAs) profiles across the entire cohort and within *TP53*-mutated cases (n = 64), allowing the identification of events specifically associated with the mutation (**Figure 1C**). Notably, losses of chromosome 3p appeared as one of the most frequent large-scale deletions in *TP53*-mutated cases (16/64, 25% of samples with available WGS), alongside known associated hotspots on 5q, 7q, and 17p.

### Recurrent deletions of chromosome 3p co-occur with -5/del(5q) and *TP53* alterations

To specifically characterize -3/del(3p) AML, all Leucegene specimens presenting deletions of the chromosome (whole chromosome 3, n = 4; or limited to the small arm, n = 14) were subjected to further analysis (**Methods and Sup. Table S2** for detailed information about the cohort). Although the deletion often involved a larger segment, the distribution of the median log2 copy number ratio along chromosome 3 revealed a common deleted region (CDR) spanning the 3p22.1 to 3p13 cytobands (global minimum, **Figure 2A**).

**Figure 2.**
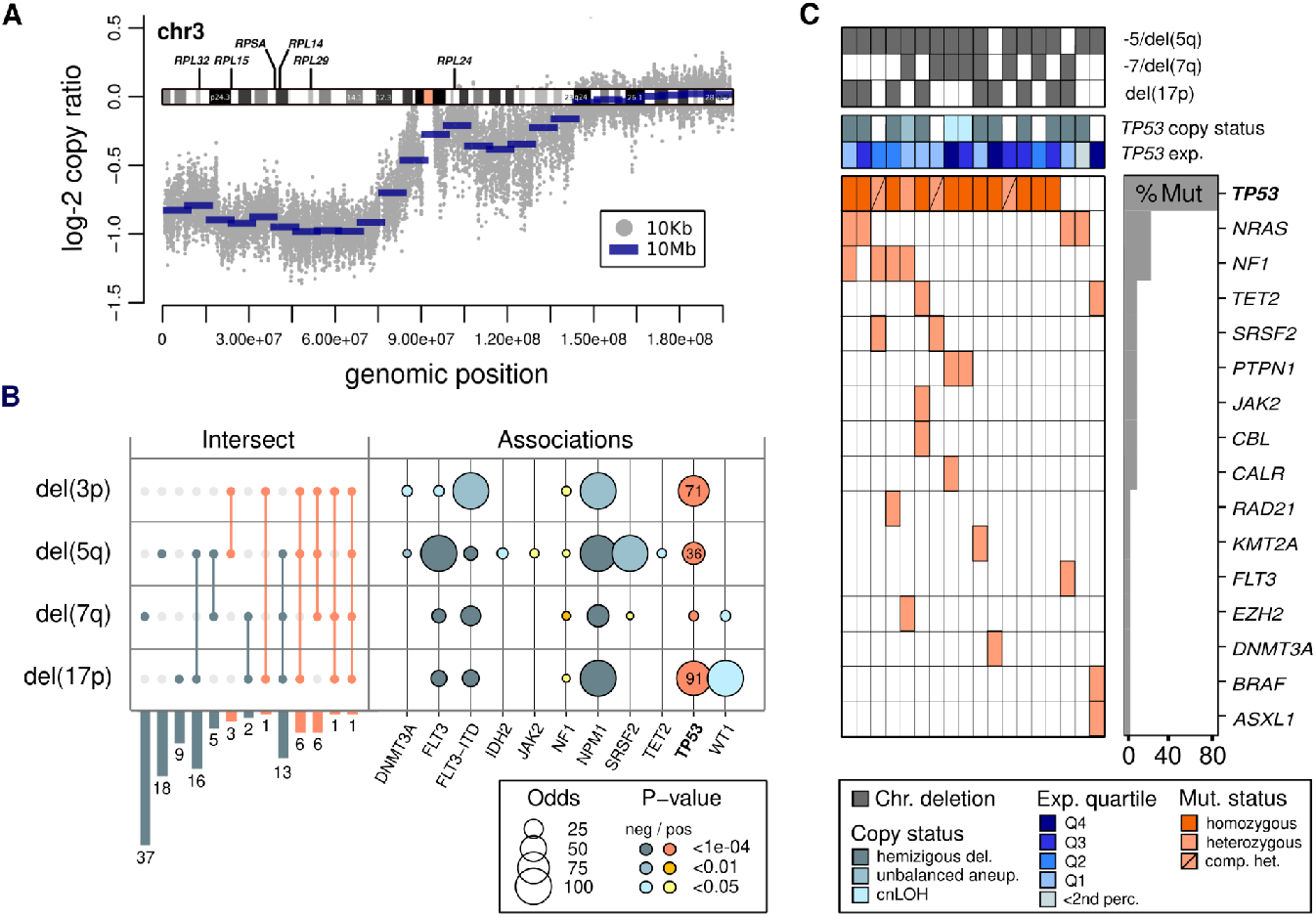
**A**. Chromosome 3p deletion footprint of -3/del(3p) specimens obtained by plotting median log2 copy ratio calculated for windows of 10Kb (depicted by gray dot) along chromosome 3. Genomic positions are indicated along the x-axis and the schematic representation of chromosome 3 is drawn at a log2 copy ratio value of 0 corresponding to a normal diploid state. Blue solid segments represent the median log2 copy ratio calculated for overlapping windows of 10 Mb. **B. Left panel**: Upset plot representation of the intersect between specimens presenting large 3p, 5q, 7q or 17p deletions. Orange lines and bars depict intersects implying the del(3p) group. **Right panel**: Bubble plot representation of associations between specimens 3p, 5q, 7q or 17p deletions and small mutations. -3/del(3p) specimens are excluded from other tested groups; for each tested group, remaining AML specimens of the Leucegene cohort are used as control group. Bubble sizes and colors are representative of association odds and p-values, respectively. Only variables presenting a significant association (positive or negative) with one of the groups are represented. **C**. Mutation heatmap of -3/del(3p) AML. Genes (y-axis) composing the heatmap were either mutated in one or more -3/del(3p) specimens (x-axis). Genes were ordered (from top to bottom) based on their mutation frequencies (indicated by the gray bar graph on the right). Specimens were grouped according to their mutation status (from left to right). Co-occurring large deletions of chromosome 5, 7 and 17, *TP53* copy status and expression quartiles are indicated in 2 panels at the top of the figure. aneup.: aneuploidy, comp. het.: compound heterozygous, cnLOH: copy neutral loss of heterozygosity, perc.: percentile. See legend at the bottom for the color code.

The resulting group (n = 18) was essentially composed of AML with myelodysplasia-related changes (n = 16/18, p < 1e-04, **Table 1**), lower white blood cell count than other AML (p < 5e-03, **Table 1**) and an association with erythroid leukemia M6 morphology under the French-American-British (FAB) classification (p < 5e-03, **Table 1**). All -3/del(3p) specimens harbored a complex karyotype (CK) and were associated with adverse cytogenetic risk (n = 18/18, p << 1e-04, **Table 1**). Most presented -5/del(5q) alterations (n = 16/18, *vs*. other AML: p << 1e-04, *vs*. CK: p < 0.05), suggesting cooperative losses, and often carried additional -17p and -7/del(7q) (n = 9 and 8/18, respectively) (**Table 1, Figure 2B and 2C**).

**Table 1.** Characteristics of the del(3p) AML cohort.

|  |  | del(3p)<br>(n=18) | other<br>(n=638) | Assoc. type with<br>del(3p) | p-values |
| --- | --- | --- | --- | --- | --- |
| Age (median, range) |  | 58.5 (39-82) | 57 (17-87) | - | - |
| Gender | Female | 5 | 269 | - | - |
|  | Male | 13 | 369 | - | - |
| WBC (median, range) |  | 8.9 (1.5-128.1) | 28.5 (0.7-447.0) | - | **<5e-03** |
| Cytogenetic risk | Undetermined | 0 | 2 | - | - |
|  | Adverse | 18 | 154 | pos | **<<1e-04** |
|  | Intermediate | 0 | 402 | neg | **<1e-04** |
|  | Favorable | 0 | 80 | - | - |
| FAB | M0 | 2 | 40 | - | - |
|  | M1 | 3 | 157 | - | - |
|  | M2 | 2 | 96 | - | - |
|  | M3 | 0 | 30 | - | - |
|  | M4 | 1 | 104 | - | - |
|  | M5 | 0 | 87 | - | - |
|  | M6 | 3 | 7 | pos | **<5e-03** |
|  | M7 | 0 | 3 | - | - |
|  | NC/other | 7 | 114 | - | - |
| WHO<br>classification<br>2016 | AML-MRC | 16 | 175 | pos | **<1e-04** |
| Subgroup | -5/del(5q) | 16 | 52 | pos | **<<1e-04** |

AML-MRC, acute myeloid leukemia with myelodysplasia-related changes; CK, complex karyotype; FAB, French-American-British classification; WBC, white blood cell count; WHO, World Health Organization

Expectedly, *TP53* was the most frequently mutated gene (n = 15/18, **Figure 2B and 2C, Sup. Table S2**) and presented a strong enrichment in this subgroup when compared to other AML (p << 1e-04; **Figure 2C**). Interestingly, odds to identify *TP53* mutations were higher for AML with -3/del(3p) alterations than for specimens of the -5/del(5q) group (71x *vs*. 36x), suggesting a strong cooperative effect between p53 loss-of-function and deletions of chromosome 3p (**Figure 2B**). As previously observed^10,11^, due to co-occurring hemizygous deletions or copy neutral loss of heterozygosity (cnLOH), most of the patients presented a homozygous expression of the *TP53* mutated allele (n = 11/18, **Figure 2C and Sup. Table S2**). Finally, three cases harbored compound heterozygous mutations (**Figure 2C**). Overall, only one out of the 18 -3/(del3p) cases presented no identifiable alteration of *TP53* (including small mutations, copy alterations or aberrant expression).

While available data did not allow concluding on the acquisition timeline of main co-occurring alterations, copy number ratios indicated that 3p deletions were clonal, displaying values consistent with those of 5q events and variant allele frequencies (VAFs) of *TP53* mutations (**Sup. Figure S2**). Of note, patients in the -3/del(3p) had very poor overall survival with no significant difference compared to patients with -5/del(5q) without alteration of chromosome 3p (**Sup. Figure S3**).

### -3/del(3p) alteration is associated with a global reduction in RPGs expression

A differential expression analysis comparing the del(3p) subgroup (n = 18) to other AML (n = 638) identified a general downregulation of cytosolic RPGs (n = 99, logFC < 0) with 65 RPGs showing a significant expression drop (n = 65, logFC < -0.5, FDR < 0.05, **Figure 3A and Sup. Table S3**). As expected, RPGs located on chromosome 3p (**Figure 2A**) were among the most consistently differentially expressed genes (DEGs) in -3/(del3p) AML. These included *RPL15* (3p24.2, top 1 downexpressed gene) and *RPL14* (3p22.1, top 3), both associated with the MDS- and AML-predisposing syndrome Diamond-Blackfan anemia when mutated^12-14^, a blood disorder characterized by a selective reduction of erythroid precursors and progenitors provoked by ribosomopathies. Additionally, the 5q encoded MDS-related haploinsufficient driver *RPS14* (5q33.1)^15^ and *RPS23* (5q14.2) were also among the top DEGs.

**Figure 3.**
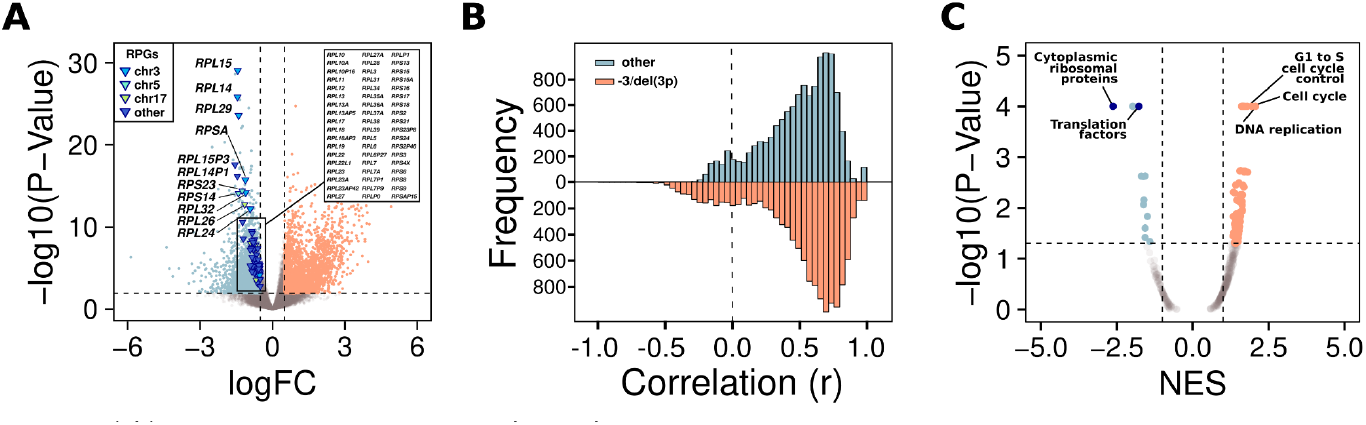
**A**. Volcano plot representation of the differential expression analysis conducted on RNAseq data comparing del(3p) AML *versus* other AML of the Leucegene cohort. The horizontal dashed line indicates an adjusted p-value of 0.01 and vertical dashed lines indicate log fold change (logFC) of 0.5 and -0.5. Orange and blue dots correspond to genes significantly over- and under-expressed (logFC > |0.5|, FDR < 0.01), respectively. Triangles depict differentially expressed RPGs (color code indicates the location: blue, chromosome 3; turquoise, chromosome 5; green, chromosome 17; and dark blue, other chromosomes). **B**. Density histogram representation of the pairwise correlation of cytosolic RPGs. **C**. Volcano plot representation of enrichments obtained from the gene set enrichment analysis (GSEA) comparing -3/del(3p) AML to other AML and using Wikipathways as gene set. NES: normalized enrichment score. Significant hits with positive or negative NES are depicted as orange and blue dots, respectively.

To test for an eventual perturbation of the co-regulation network of RPGs in this context, we conducted a pairwise correlation analysis of cytosolic ribosomes (**Figure 3B and Sup. Figure S4**). No major discoordination of expression in -3/del(3p) AML was observed compared to other AML and most genes remained highly correlated. Furthermore, genes carried by chromosome 3p such as *RPL15*, presenting a consistent expression drop in -3/del(3p) AML, maintained a comparable correlation level than observed in other AMLs (**Sup. Figure S5**).

A gene set enrichment analysis (GSEA) expectedly confirmed the significant downregulation of genes involved in ribosomal production and translation pathways and, interestingly, showed a strong positive enrichment of genes associated with DNA replication and cell cycle (**Figure 3C**). Overall, these results suggest the presence of a somatic ribosomopathy combined with a paradoxical active proliferation.

### Combination of chromosome 3p and 5q RPG-CNAs with mutated *TP53* is associated with the weakest overall expression of cytosolic RPGs in AML

To obtain a unique and representative expression value per specimen for all the cytosolic RPGs together, we computed a ribosomal eigengene vector (first principal component of a PCA of the RPGs, **Methods and Sup. Table S2**). Total protein synthesis was assessed by flow cytometry using puromycin labeling (**Methods**) and confirmed that the calculated eigengene expression was indeed representative of the translational levels in tested cells (**Figure 4A and Sup. Figure S6**). This value was therefore used as the reference variable to test associations and correlations throughout subsequent analyses (see **Table 2** for associations with clinical variables). Low ribosomal eigengene values were notably associated with an adverse cytogenetic risk and enriched in FAB M5 and M6 morphologic subtypes, whereas high ribosomal expressions were associated with M0 and M1 leukemias. Testing the distribution of this eigengene vector in groups defined by recurrent genetic anomalies identified in the Leucegene cohort (including structural variants and small mutations; n = 70 variables identified in ≥ 5 specimens), distinguished -3/del(3p) and, to a lesser extent, -5/del(5q) alterations from other anomalies (**Figure 4B**). Note that del(5q) AML showed much less significance when del(3p) and/or *TP53*m co-occurrences were excluded (see del(5q)* and del(5q)^‡^ in **Figure 4B**).

**Table 2.** Ribosomal eigengene expression *versus* clinical characteristics.

|  |  | Low RPG eigengene (tier1, n=215) | High RPG eigengene (tier3, n=222) | Assoc. type with low RPG eigengene | p-values |
| --- | --- | --- | --- | --- | --- |
| **Age** (median, range) |  | 58 (17-87) | 57 (18-84) | - | - |
| **Gender** | Female | 89 | 90 | - | - |
|  | Male | 126 | 132 | - | - |
| **WBC** (median, range) |  | 28.9 (0.7-327.0) | 20.5 (0.9-447.0) | - | - |
| **Cytogenetic risk** | Undetermined | 0 | 1 | - | - |
|  | Adverse | 76 | 56 | pos | **<0.05** |
|  | Intermediate | 24 | 25 | - | - |
|  | Favorable | 115 | 140 | - | - |
| **FAB** | M0 | 8 | 25 | neg | **<0.01** |
|  | M1 | 25 | 82 | neg | **<<1e-04** |
|  | M2 | 23 | 37 | - | - |
|  | M3 | 14 | 5 | pos | **<0.05** |
|  | M4 | 37 | 21 | pos | **<0.01** |
|  | M5 | 51 | 13 | pos | **<1e-04** |
|  | M6 | 9 | 0 | pos | **<5e-03** |
|  | M7 | 2 | 1 | - | - |
|  | NC/other | 46 | 38 | - | - |
| **Mutation** | *TP53* | 44/204* | 14/209* | pos | **<1e-04** |

FAB, French-American-British classification; WBC, white blood cell count; \*documented

**Figure 4.**
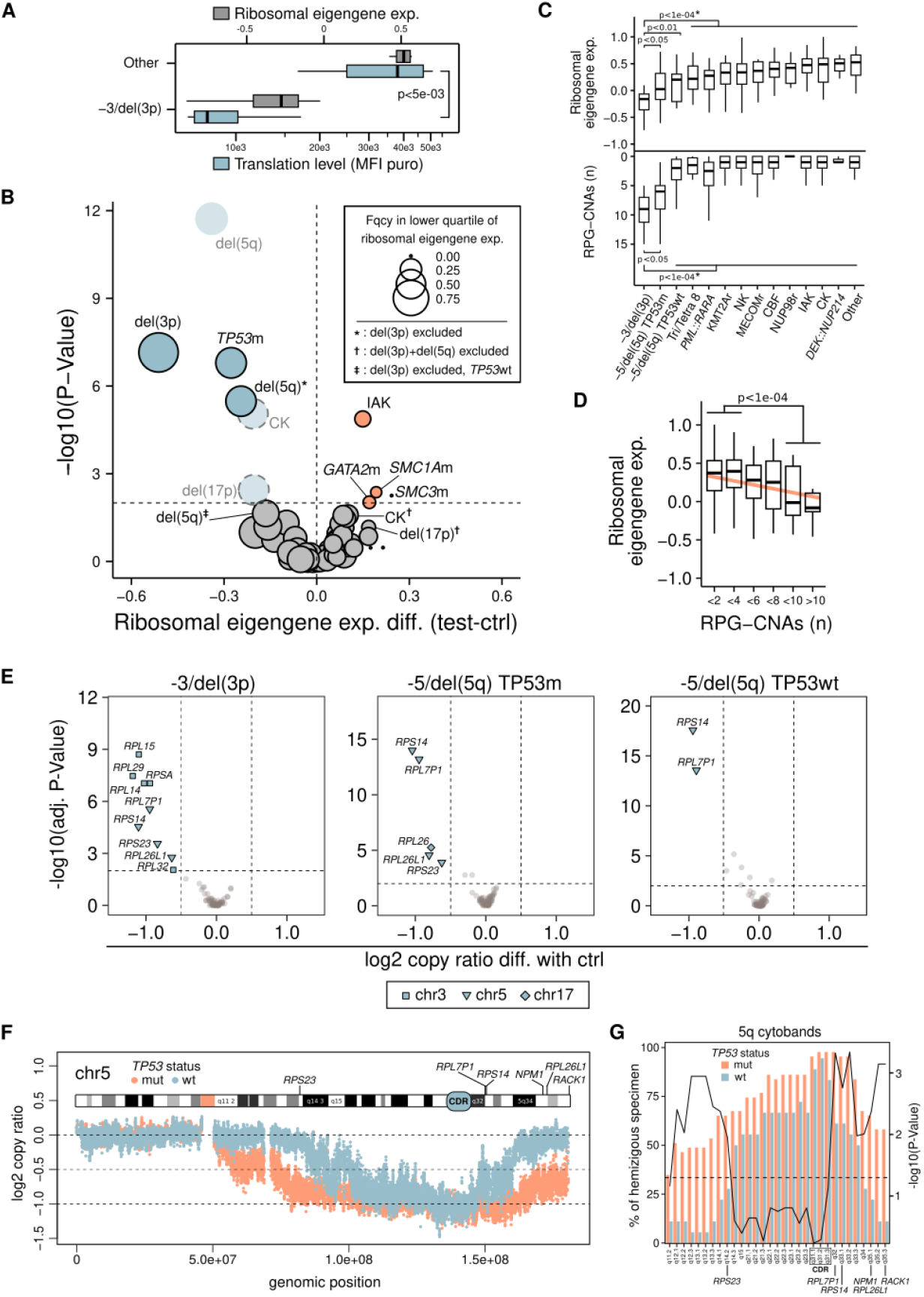
**A**. Translational levels (assessed by puromycin labeling, blue boxes, bottom x-axis) of a selection of -3/del(3p) (n = 6) and other AML (n = 6), presenting low and high ribosomal eigengene values respectively (gray boxes, top x-axis). The p-value resulting from the comparison between the 2 groups is directly indicated on the figure. MFI, mean fluorescence intensity. **B**. Volcano plot representation resulting from the comparison of ribosomal eigengene distributions presented by test groups – each being defined by a recurrent genetic anomaly identified in the Leucegene cohort, including structural variants and small mutations (n = 70 variables identified in ≥ 5 specimens) – and controls (ctrl), composed of other AML not presenting the considered variable (see legend in the top right corner of the figure for specific exclusions in tested groups). The x-axis and y-axis represent the differences of mean ribosomal eigengene expression and p-values resulting from Wilcoxon tests comparing the eigengene distributions between test and control groups, respectively. Bubble sizes are representative of the frequency of specimens in each tested group with a ribosomal expression located in the lower quartile of the eigengene vector (see legend in the top right corner of the figure for scale). CK: complex karyotype, IAK: intermediate abnormal karyotype (except isolated trisomy/tetrasomy 8). **C**. Ribosomal eigengene expression values (**top panel**) and number of RPG-CNAs (**bottom panel**, inverted axis) according to -3/del(3p) specimens and other AML of the Leucegene cohort (groups are indicated along x-axis). P-values resulting from the tests comparing each group to the -3/del(3p) AML are directly indicated on the figure (* p-value obtained for the comparison between the test and each control group under the transversal bar). CBF: core-binding factor AML including *RUNX1*::*RUNX1T1* and inv(16)(p13.1q22), CK: complex karyotype, NK: normal karyotype, IAK: intermediate abnormal karyotype (except isolated trisomy/tetrasomy 8). **D**. Ribosomal eigengene expression values according to the number of RPG-CNAs (mutually exclusive groups, the solid orange line results from a least squares regression). The p-value resulting from the comparison between extreme groups (RPG-CNAs <4 *vs*. >8) is directly indicated on the figure. **E**. Volcano plot representation of the log2 copy ratio analysis of RPGs comparing -3/del(3p) specimens *vs*. other AML (**left panel**), -5/del(5q) specimens mutated for *TP53* (TP53m) *vs*. other AML (**central panel**), and -5/del(5q) specimens wild-type for *TP53* (TP53wt) *vs*. other AML (**right panel**). -3/del(3p) and -5/del(5q) subgroups are mutually exclusive. The x-axis represents the log2 copy ratio difference between the 2 groups. The horizontal dashed line indicates a FDR of 0.01 and vertical dashed lines indicate log2 copy ratio differences of 0.5 and -0.5. Blue dots correspond to RPGs presenting a significant loss of copy (log2 copy ratio difference < -0.5, FDR < 0.01). Significant RPGs located on chromosome 3, 5 and 17 are depicted by blue squares, triangles and diamonds, respectively, while other RPGs are indicated by gray dots. **F**. Chromosome 5q deletion footprint of wild-type and mutated -5/del(5q) specimens obtained by plotting median log2 copy ratio calculated per group for windows of 10Kb along chromosome 5. Windows are depicted by blue and orange dots for wild-type and mutated -5/del(5q) specimens, respectively. Genomic positions are indicated along the x-axis and the schematic representation of chromosome 5 is arbitrarily drawn at a log2 copy ratio value of 0.5. The common deleted region (CDR) of del(5q) specimens is directly indicated on the representation of chromosome 5. **G**. Barplot representation of the percentage of hemizygous *TP53*-mutated (orange bars) and wild-type (blue bars) -5/del(5q) specimens for 5q cytobands (left axis). The black curve indicates the p-values resulting from Fisher’s exact tests comparing *TP53*-mutated to wild-type -5/del(5q) AML for each cytoband (right axis). The horizontal dashed line indicates a p-value of 0.05.

Overall, the eigengene followed a linear trend with lowest values associated with the combined low expression of RPGs located on both chromosome 3 and 5 (**Sup. Figure S7**), such as *RPL15* (3p24.2) and *RPS14* (5q33.1) – followed by (in ascending order) *RPL15*^low^/*RPS14*^high^, *RPL15*^high^/*RPS14*^low^ and *RPL15*^high^/*RPS14*^high^. Expectedly, -3/del(3p) cases presented significantly lower eigengene values compared to other AML subgroups (**Figure 4C, top**) and was directly followed by -5/del(5q) AML mutated for *TP53* with a lower ribosomal gene expression than non-mutated specimens, stressing the importance of p53 loss-of-function in a ribosomopathic context.

Importantly, in the Leucegene cohort, the number of deleted cytosolic RPGs per specimen was found to correlate with the general ribosomal eigengene **(Figure 4D**). This correlation indicates a quantitative relationship between the RPG-CNA burden and the global expression of the RPG network, which cannot be explained solely by the downregulation of deleted genes (**Sup. Figure S8**).

In line with these results, -3/del(3p) and -5/del(5q) AML mutated for *TP53* presented the highest number of RPG-CNAs in the cohort with a median of 9 and 6 events per specimen, respectively, while non-mutated -5/del(5q) cases only presented a median value of 2 deletion events (**Figure 4C, bottom**). Overall, despite few discrepancies such as *PML::RARA* positive leukemias presenting a higher expression than suggested by the number of RPG losses, most subgroups showed a correlated relationship between RPG-CNAs and RPG eigengene.

Recalling the co-occurrence of copy losses in the -3/del(3p) cohort, chromosomes 3, 5 and 17 had the higher frequencies of RPG-CNAs in the subgroup, with recurrently and significantly deleted RPGs restricted to chromosomes 3p and 5q (**Figure 4E**). Although chromosome 19 was the first contributor at the scale of the whole Leucegene cohort given its high density in RPGs (∼0.25 RPGs/Mb *vs*. < 0.1 for other chromosomes), it showed no enrichment in the -3/del(3p) specimens (**Sup. Figure S9**). As for -5/del(5q) AML, both *TP53* mutated and wild-type cases presented chromosomes 5 and 17 as top contributing chromosomes but with significantly stronger enrichments when mutated (**Sup. Figure S9**).

Interestingly, we and others previously identified a relationship between longer 5q deletions and alterations of *TP53*^11,16,17^ – suggesting a cooperation between the copy-loss of genes located in the area of chromosome 5 specifically associated with inactivation of p53. Accordingly, the comparison of 5q deletion footprints depending on *TP53* status revealed that although sharing the CDR, wild-type -5/del(5q) specimens usually present events with boundaries limited from 5q14.3 to 5q33.3. In contrast, deletions in mutated cases more often extend from 5q12.1 to the chromosome end (**Figure 4F and 4G**), hence frequently include *RPS23* (5q14.2) and *RPL26L1* (5q35.1) – 65% and 22% of mutated and wild-type cases, respectively, are hemizygous for 5q14.2 and 5q35.1, (p-value < 5e-03, **Figure 4G**) – which explains the specific association of their copy-losses with mutated cases (**Figure 4E**). Furthermore, as reported here and in our recent study on -5/del(5q)^11^, most of these mutated specimens present a homozygous expression of the mutated allele due to large or focal deletions of chromosome 17p which translated here as a significant association of *RPL26* deletion located at 17p13.1 close to *TP53* (**Figure 4E**). Note that no significant difference in 5q deletion footprint was identified between *TP53*-mutated -3/del(3p) and -5/del(5q) specimens (**Sup. Figure S10**). Overall, these data demonstrated the importance of the combination of events on chromosome 3p, 5q (including *TP53*-dependent differentially deleted regions) and 17p (including *TP53* mutations) to obtain the stronger ribosomopathy-like phenotype, as observed in most -3/del(3p) specimens.

### *TP53*-dependent events could contribute to the destabilization of ribosomal network in AML

To evaluate the coregulation of key biological processes and RPGs in AML, we assessed the correlation between the ribosomal eigengene vector and genes constituting the 50 hallmark sets from the Human Molecular Signatures Database (MSigDB) (**Sup. Figure S11A and Sup. Table S4**). This analysis revealed that low RPGs expression is rather associated with a proinflammatory state, sets such as “complement”, “inflammatory response”, “TNF-α signaling via NF-κB” and “interferon-γ response” being among the top negatively correlated pathways. Although this may result from the general *TP53*-mutated context of low eigengene expressors, cellular stress and impairment of the ribosome-dependent inhibition of the inflammatory response are also likely at play^18,19^. On the other hand, gene sets dependent on Myc (“MYC targets” V1 and V2), one of the major modulators of ribosome biogenesis^19^, presented the strongest positive correlation with the eigengene (**Figure 5A and Sup. Figure S11A**), while mTOR signaling, the other master regulator^19^, showed no evidence of co-modulation (**Sup. Figure S11A**).

**Figure 5.**
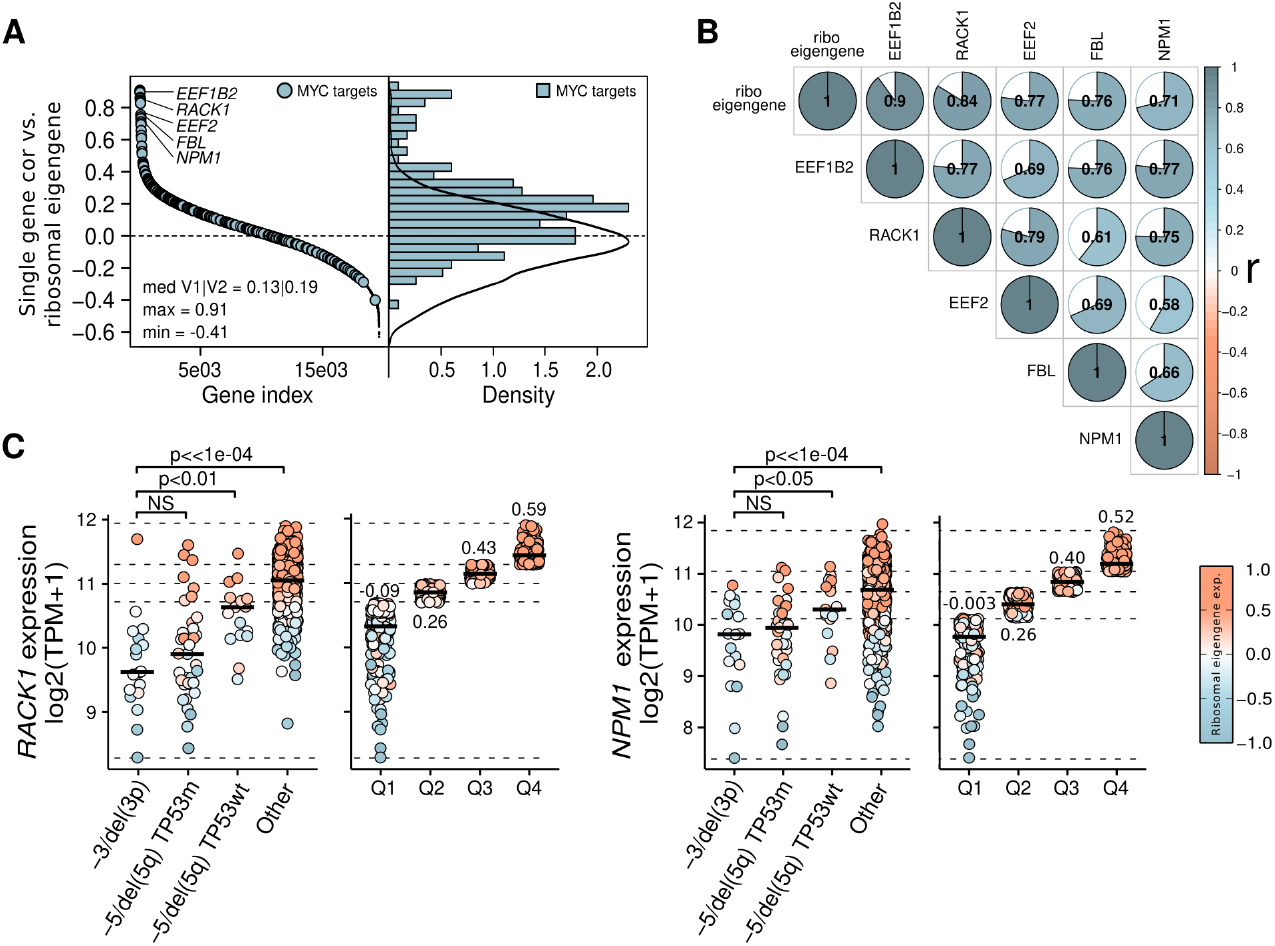
**A**. Single gene correlation versus ribosomal eigengene (Pearson’s correlation, **left panel**) and corresponding density plot (**right panel**). MYC targets (V1 and V2 combined from the hallmark set of the Human Molecular Signatures Database, MSigDB) are indicated by blue dots (left) or bars (right). Median (med), maximum (max) and minimum (min) correlation values for MYC targets are directly indicated on the left panel. The solid black curve in the right panel corresponds to the density obtained from the 48 other hallmark sets together. **B**. Pairwise correlation plot comparing a selection of MYC targets and the ribosomal eigengene expression in the Leucegene cohort (n = 656 AML). Area and blue shades of colored surfaces in pie charts (color scale indicated at the right of the panel) are representative of Pearson’s correlation coefficients (r, also indicated in each chart). **C**. *RACK1* and *NPM1* expression in the -3/del(3p), -5/del(5q) TP53m or TP53wt and other AML of the Leucegene cohort (**left panel of each plot**) or distributed into expression quartile subgroups (**right panel of each plot**). Median values are indicated by black lines on each dotplot. As indicated in the legend (right of the figure), the color code is representative of the ribosomal eigengene expression. P-values resulting from the comparison between groups are indicated on the plot (NS, non-significant). Transversal dotted lines delimit quartiles. Median ribosomal eigengene expression values for each expression quartile are directly indicated on the plots.

Running a GSEA comparing the -3/del(3p) subgroup to other AML specimens also confirmed the conjoined downregulation of ribosomes and MYC targets (**Sup. Figure S11B**). More specifically, apart from several RPGs expectedly coregulated (**Sup. Table S4**), other genes potentially contributing to the ribosomopathic phenotype were found among the top correlated MYC targets (**Figure 5B**): the two elongation factors EEF1B2 and EEF2, the fibrillarin (FBL) that promotes early pre-rRNA processing^20^ (r = 0.90, 0.77, 0.76, respectively, **Sup. Table S4 and Sup. Figure S11A**), and particularly interesting given their role and their location in the *TP53*-related differentially deleted region at the end of chromosome 5 (5q35.3 and 5q35.1, **Figure 4F and 4G**), the Receptor for Activated C Kinase 1 (RACK1) and the nucleophosmin 1 (NPM1) (r = 0.84 and 0.71, respectively, **Sup. Table S3, Sup. Table S4 and Sup. Figure S11A**). RACK1 is involved in translation and ribosome quality control^21-28^, as for NPM1, besides its importance in AML, the dysregulation represents a particular interest here given its role in different phases of ribosome synthesis and nucleolar stress response^29-33^. Detailing *RACK1* and *NPM1* expression profile in the whole Leucegene cohort confirmed that only low expressors reached low levels of ribosomal eigengene (**Figure 5C**), with -3/del(3p) and *TP53*-mutated -5/del(5q) AML – which present highest frequencies of *RACK1* and *NPM1* deletion (**Figure 4G**) – showing the lowest expression levels (**Figure 5C, Sup. Table S3**). Expectedly, specimen 09H045 with the highest ribosomal eigengene expression of the -3/del(3p) subgroup and no alteration of chromosome 5 (**Sup. Table S2**), also presented the highest expression of both genes in the group and was the only -3/del(3p) specimen in the top expression quartiles of the Leucegene cohort (**Figure 5C**). Note that, while it has been hypothesized that a reduction of NPM1 dosage – as provoked by commonly encountered small insertions in the last exon of the genes (NPM1c) – could alter ribosome synthesis^34^, NPM1c alone was not associated with a variation of ribosomal expression (n_*NPM1*wt_ and n_*NPM1*c_ > 400 and 200, respectively, **Sup. Figure S11C**).

### Reduction of ribosomal protein levels provoked by HSP90 inhibitors lead to the sensitization of low RPG expressors

Based on these findings, we hypothesized that the transcriptional and translational specificities associated with ribosomopathy-like profiles could represent a therapeutic vulnerability. To explore this, we utilized data from a pre-existing single-dose high-throughput screening (HTS) assay, which tested over 10,000 structurally diverse compounds across 56 primary AML specimens from the Leucegene cohort (**Sup. Figure S12**). Compounds exhibiting similar inhibitory profiles across specimens – potentially targeting the same cellular pathways – were grouped into Compound Correlation Clusters (CCCs) using hierarchical clustering based on a minimum spanning tree approach (**Figure 6A, Methods**).

**Figure 6.**
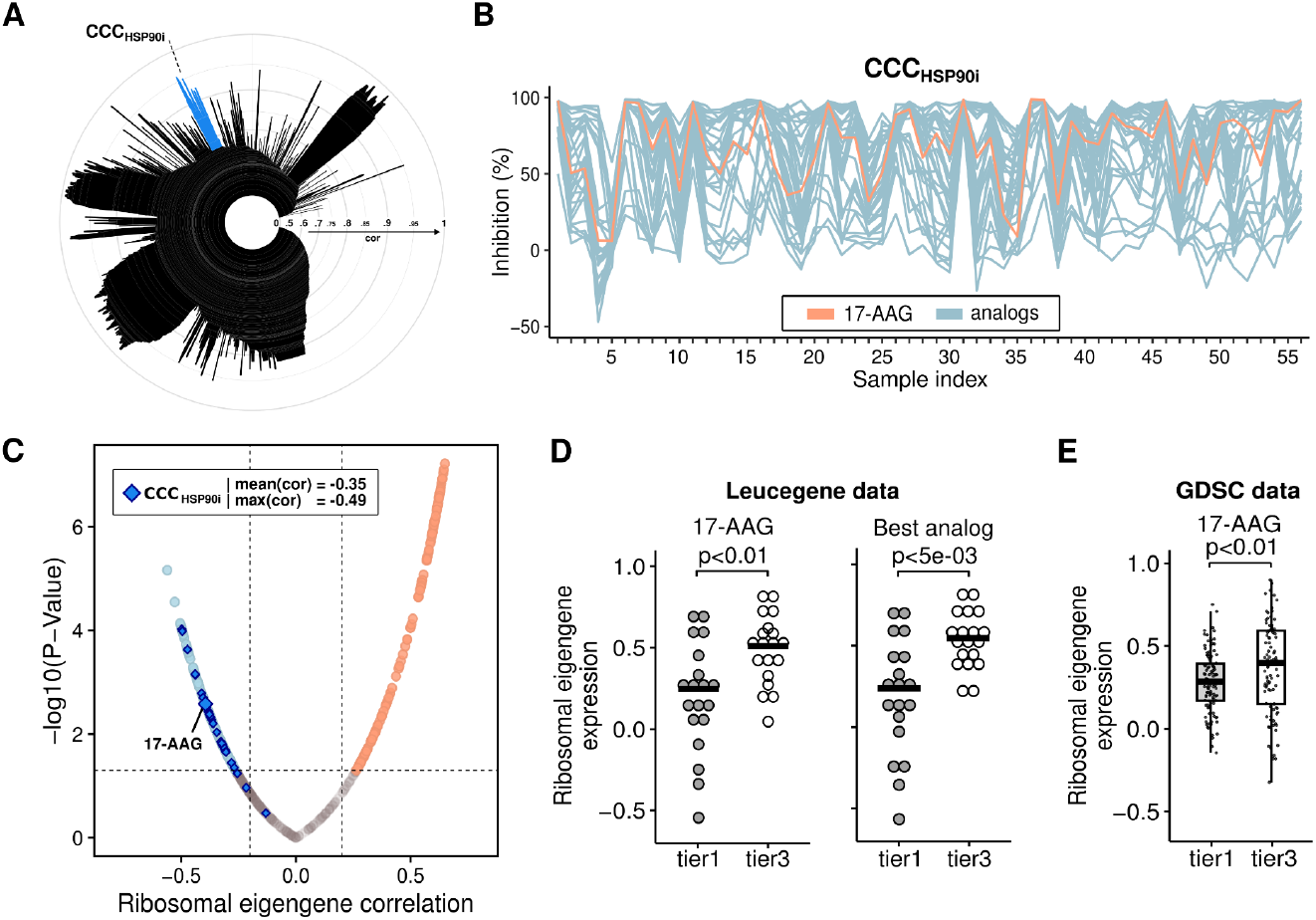
**A**. Icicle representation of selective hit compounds of the initial single-dose high-throughput screening (HTS) assay depicting peaks of highly correlating inhibitory profiles constituting Compound Correlation Clusters (CCCs). Concentric circles depict levels of correlation as indicated on the figure (from 0 at the center to 1 at the edge). CCC_HSP90i_ composed of the 17-AAG and 33 of its analogs is depicted by a blue peak. **B**. Inhibition footprint (%) obtained for compounds composing the CCC_HSP90i_. Orange and blue lines represent data corresponding to the 17-AAG and its analogs, respectively. **C**. Volcano plot representation of the correlation level between compounds constituting the CCCs (Compound Correlation Clusters) and the ribosomal eigengene expression values obtained for tested specimens. The horizontal dashed line indicates a p-value of 0.05 and vertical dashed lines indicate correlation of 0.2 and -0.2. Orange and blue dots correspond to compounds significantly correlated and inversely correlated (correlation > |0.2|, p-value < 0.05), respectively. Dark blue diamonds depict compounds from the CCC_HSP90i_ composed of the 17-AAG (indicated by a larger blue diamond) and 33 of its analogs. **D**. Ribosomal eigengene expression values according to tier 1 (“sensitive” tier) and tier 3 (“resistant” tier) sensitivity groups for 17-AAG (**left panel**) and the best analog (**right panel**), i.e. presenting the strongest difference between the 2 groups. **E**. Ribosomal eigengene expression values according to tier 1 (“sensitive” tier, n = 114 cell lines) and tier 3 (“resistant” tier, n = 98) 17-AAG sensitivity groups determined using data obtained from The Genomics of Drug Sensitivity in Cancer (GDSC) database.

We then evaluated the correlation between RPG eigengene values, calculated for each tested specimen, and the inhibitory profiles of each identified CCC. This analysis highlighted a chemical cluster centered on 17-AAG/Tanespimycin, a heat shock protein 90 (HSP90) chaperone inhibitor (HSP90i) that has been investigated in multiple clinical trials for various cancer types^35-39^, along with 33 of its analogs. Notably, this cluster showed the strongest and most consistent anticorrelation with the RPG eigengene, with 31 out of 34 compounds in the CCC displaying significant anticorrelation (**Figure 6A-D**). These findings suggest an association between low ribosomal expression and sensitivity to HSP90 inhibition.

To validate this observation, we leveraged The Genomics of Drug Sensitivity in Cancer (GDSC) database^40^ as an independent dataset. Despite the substantial genetic variability among the >200 cancer cell lines tested, the association between ribosomal expression and HSP90 inhibitor sensitivity was confirmed (**Figure 6E**).

We further conducted a targeted chemical screen for validation, testing three HSP90 inhibitors – 17-AAG, Geldanamycin, and Alvespimycin – on a cohort of -3/del(3p) AML samples (n = 13) and control specimens (n = 25) with varying ribosomal expression profiles (**Figure 7A; Sup. Table S5**). The three compounds exhibited similar response profiles, with the strongest correlation observed between 17-AAG and Alvespimycin (Pearson’s coefficient = 0.79, **Figure 7B**). -3/del(3p) specimens showed a significantly increased sensitivity to HSP90 inhibition compared to controls, independently of p53 status alone (**Figure 7A and 7C**). Overall, although this smaller screen did not yield a statistically significant association between ribosomal expression and sensitivity, these results remained consistent with the initial findings (**Figure 7C**).

**Figure 7.**
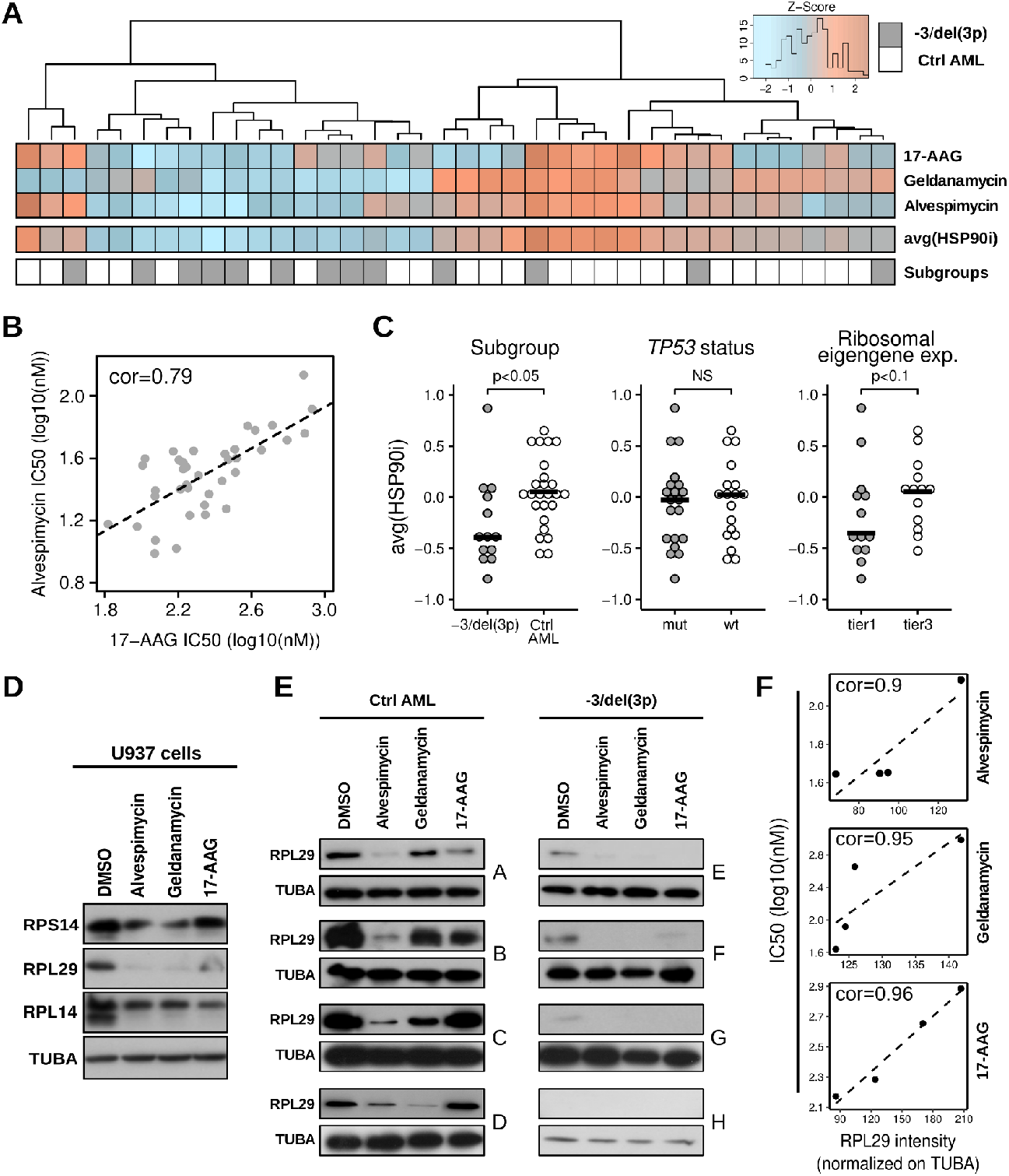
**A**. Sensitivity heatmap for HSP90i (17-AAG, Geldanamycin and Alvespimycin). Color code is representative of Z-scores (see color key at the top) calculated for each compound from IC50 values. Color code in the bottom line indicates specimen subgroups (−3/del(3p) or ctrl AML). The tree at the top depicts the unsupervised hierarchical clustering computed from IC50 values. **B**. Correlation plot comparing responses to 17-AAG and Alvespimycin (Pearson’s correlation coefficient r = 0.79). The dotted line results from a least squares regression. **C**. Response to HSP90 inhibitors for -3/del(3p) specimens *vs*. control AML (**left panel**), according to TP53 status (**central panel**) and ribosomal eigengene expression (**right panel**, tier 1 = low expressors and tier 3 = high expressors). The average response to HSP90 inhibitors (avg(HSP90i)) corresponds to the mean value of rescaled IC50 vectors (from -1 to 1) obtained for each inhibitor. P-values resulting from the comparison between groups or conditions are indicated on the plot. **D**. Representative blots showing RPS14, RPL14, RPL29 and tubulin alpha (TUBA, loading control) levels as assessed by western blot analysis of total proteins extracted from U937 cells exposed to DMSO or HSP90 inhibitors (Alvespimicin, Geldanamycin and 17-AAG (1M)) for 24hrs. **E**. Representative blots showing RPL29 and tubulin alpha (TUBA, loading control) levels as assessed by western blot analysis of total proteins extracted from primary AML cells (samples A to D: control AML without -3/del(3p) alteration; samples E to H: -3/del(3p) AML) exposed to DMSO or HSP90 inhibitors (Alvespimicin, Geldanamycin and 17-AAG (1M)) for 24hrs. **F**. Correlation plots comparing RPL29 intensity (normalized on tubulin alpha, TUBA) with responses to HSP90 inhibitors. The dotted lines result from least squares regressions. The Pearson’s correlation coefficient calculated for each compound is directly indicated on the plots (top left).

Finally, drawing on prior reports indicating that HSP90 inhibition or knockout leads to ribosome degradation or reduced ribosomal protein levels^41,42^, we investigated whether this mechanism could explain the observed sensitization of AML with low RPG expression. Western blot analysis of three ribosomal proteins (RPS14, RPL14, and RPL29) in U937 cells treated with 17-AAG, Geldanamycin, and Alvespimycin revealed a marked reduction in ribosomal protein levels across all three HSP90 inhibitors (**Figure 7D**). Complementarily, western blotting of RPL29 in primary AML cells with (n = 4) and without (ctrl AML, n = 4) del(3p) alterations was performed following treatment with the three HSP90 inhibitors (**Figure 7E**). In control AML specimens, a reduction in RPL29 levels was observed following treatment. Furthermore, these levels were found to correlate with the IC50 values (r ≥ 0.9, **Figure 7F**), indicating that the sensitivity to these inhibitors is likely dependent on RPG abundances. In del(3p) AML cells – which already exhibited markedly low baseline RPL29 levels – treatment with HSP90 inhibitors resulted in a further reduction, decreasing RPL29 to undetectable or virtually undetectable levels (**Figure 7E**).

These findings highlight the significant ribosomal protein deficiency in -3/del(3p) AML and provide a mechanistic basis for their heightened sensitivity to HSP90 inhibition. More generally, it supports the therapeutic potential of HSP90 inhibition in ribosomopathy-like AML.

## Discussion

Although loss of chromosome 3p has been previously noted as a recurrent co-aberration with 5q deletions in AML^43^, we now provide the first specific characterization of a *TP53*-altered AML subset defined by hemizygous deletions of ribosomal protein genes (RPG) on 3p and 5q. These include *RPSA* (essential for 40S ribosomal subunit assembly and stability), *RPL29* (a 60S subunit component), *RPL14* and *RPL15* (60S subunit components and Diamond-Blackfan anemia (DBA)-associated genes), as well as the 5q-syndrome-associated haploinsufficient gene *RPS14*^15,44^. These deletions were associated with a general reduction in cytosolic RPG expression, decreased translation, creating an overall ribosomopathy-like phenotype. More broadly, our findings contribute to defining the somatic ribosomopathy in AML, suggesting it represents an underestimated driving mechanism in leukemogenesis, with the extent of ribosomal regulatory network perturbations proportional to the RPG-CNA burden. This quantitative association produces a continuum of phenotypes, with -3/del(3p) *TP53*-altered AML representing an extreme manifestation. Results also suggest that although *TP53* mutations are linked to the strongest phenotype, they are not drivers of ribosomopathy. Hence, this work highlights that *TP53*-mutated AML can be categorized as ribosomopathic or not with a full spectrum of severity that depends on the associated deletions. This observation has direct consequence for selecting AML patients in whom HSP90 inhibition should be tested, as initial studies by Carter e*t al*. suggested that epichaperome inhibition may be efficient in this context^45^.

Until recently, the consequences of disrupted ribosome biogenesis and function in cancer cells were poorly understood. However, emerging evidence suggests that these disruptions may drive the oncogenic process through multiple mechanisms^18,46^, potentially enabling a paradoxical shift from an initially hypo-proliferative state to a proliferative state^47^, matching the expression signature of del(3p) AML cells. This likely implies the selective translation of mRNAs encoding genes crucial for tumor survival and proliferation, such as oncogenes and cell cycle regulators, compounded by genomic instability – as metabolic alterations associated with ribosomal dysfunction impairs the DNA damage response – increasing mutation rates and driving the acquisition of additional oncogenic alterations^48^. Furthermore, in a leukemic context, stress resistance associated with low biosynthetic activity may confer microenvironmental advantages to pre-leukemic and leukemic clones, allowing to outcompete normal hematopoietic cells for limited resources^49^.

Interestingly, recent large-scale studies (n > 10,000 tumor genomes) revealed that more than 40% of analyzed specimens harbored hemizygous deletions of RPGs^50^, including frequent deletions of *RPSA, RPL14* and *RPL29* on chromosome 3. These results suggest that ribosomal dysregulation could play an underestimated role in cancer onset and progression. More specifically, given that tightly regulated protein synthesis is necessary for maintaining the hematopoietic stem cell (HSC) pool and allowing lineage commitment, alterations in protein synthesis rates can lead to HSCs depletion and leukemogenesis^51,52^. DBA – a syndrome caused by mutations in RPGs or ribosome biogenesis factors which critically impair erythropoiesis by compromising the high protein supply needed during erythroid maturation – illustrates how a reduction in ribosome availability leads to hematopoiesis disruption^53,54^. We show that similar mechanisms appear to operate in AML, where multiple RPG deletions seem to contribute to global ribosomal defects and are comparable to the effects of inactivating mutations^50^. Moreover, AML cases with low ribosomal protein expression showing enrichment in M6 morphologic subtype, particularly align with observations derived from studies on DBA.

As with most ribosomopathy-like tumors^50^, -3/del(3p) AML exhibited p53 loss-of-function, which allows cells to evade p53-dependent nucleolar stress responses triggered by ribosomal defects, thereby bypassing growth arrest^55,56^. Beyond this, the interaction between ribosomal defects and p53 loss-of-function takes on particular relevance in -5/del(5q) AML, as our previous work demonstrated that *TP53*-mutated AML specimens tend to harbor longer 5q deletions^11^. We now demonstrate that these extended deletions include RPG-CNAs located in differentially altered regions of the chromosome arm, likely intensifying the ribosomopathy phenotype. Together, these observations uncover novel insight into one possible cooperative oncogenic mechanism linking 5q deletions and *TP53* mutations in AML. In addition to their quantitative impact on ribosomal protein depletion, 5q deletions in *TP53*-mutated cases more frequently affect genes such as *NPM1* and *RACK1* (located at 5q35), whose downregulation could further amplify ribosomal defects. Depletion of RACK1, a core ribosomal protein of the small ribosomal subunit, has notably been associated with alteration of translational capabilities and ribosome-associated quality control^21-28^. Meanwhile, the ubiquitous role of NPM1 in rRNA transcription, maturation, and transport^29,30^, suggests its dysregulation in *TP53*-mutated specimens – showing enrichments of copy alterations – as a possible part of the cooperative set of driving events.

Beyond providing mechanistic insight, these results also reveal a potential clinical perspective for a subset of *TP53*-altered AML. While ribosomopathies in AML could modulate the expression of pro-apoptotic or drug-target proteins, potentially contributing to resistance to chemotherapy, it can also represent a lever for tumor sensitization. Through integrative chemical-genetic screening, we identified HSP90 inhibition – which exacerbates ribosomal defects by limiting ribosomal protein availability – as toxic for cells with low RPG expression, including *TP53*-altered AML with -3/del(3p).

Taken together, our data support the existence of a *TP53*-altered, ribosomopathy-like AML subtype. This not only refines our understanding of leukemogenesis in this high-risk group but also provides a rationale for exploring tailored ribosome-targeting strategies in this otherwise refractory disease.

## Material and methods

### Primary AML specimens

The Leucegene project [https://leucegene.ca/] is an initiative approved by the Research Ethics Boards of Université de Montréal and Maisonneuve-Rosemont Hospital. Leucegene AML samples (n = 691, bone marrow or blood samples) were collected from 656 patients between 2001 and 2019 and characterized by the Quebec Leukemia Cell Bank after obtaining an institutional Research Ethics Board–approved protocol with informed consent according to the Declaration of Helsinki. The Quebec Leukemia Cell Bank is a biobank certified by the Canadian Tissue Repository Network. Cytogenetic aberrations and composite karyotypes were described according to the International System for Human Cytogenomic Nomenclature 2020 guidelines^57^. Specimens were classified based on the latest recommendations from the World Health Organization (WHO) and the International Consensus Classification (ICC) of Myeloid Neoplasms and Acute Leukemias^6,58^ (unless otherwise specified). Using the analytic approach on WGS data described below, a recurrent large deletion of chromosome 3p was identified as shared by 14 AML cases. The -3/del(3p) cohort was completed to a total of 18 specimens (**Sup. Table S2**) by adding 4 cases with a complete loss of the chromosome 3 – including 2 specimens for which cytogenetics information only was available – presenting similar clinical and transcriptomic characteristics. The -5/del(5q) cohort was composed of 68 specimens (n = 52, -3/del(3p) excluded), of which 49 presented a *TP53* mutation.

### Survival analysis and Leucegene AML prognostic cohort

Survival analysis was conducted using the Leucegene AML prognostic cohort (n = 470), a subset of the Leucegene AML cohort including patients with newly diagnosed AML, excluding acute promyelocytic leukemia (APL) and patients who were treated with an intensive chemotherapy regimen with a curative intent (mostly 7+3 regimen backbone). Overall survival (OS) was evaluated using the Kaplan-Meier method. Cox proportional hazards (CPH) models were used to calculate hazard ratio (HRs) between subgroups with 95% confidence intervals (95% CI). OS was defined as time from diagnosis to time of death or last follow-up.

### Low-pass whole genome sequencing and analysis

Tumor gDNAs (n = 611) were sequenced on NovaSeq6000 S4 (paired-end 150 bp). Alignment to GRCh38 was done using the BWA aligner (v0.7.12)^59^, PCR duplicates were removed using Picard (MarkDuplicates)^60^ and a GATK (v4.1.0, BaseRecalibrator)^61^ base quality score recalibration was applied. A mean depth coverage ∼5X was reached for each sample. Identification of regions of genomic gains and losses was done using FREE-Copy number caller (v11.5, FREEC)^62^. The optimization of algorithm parameters (breakPointThreshold = 1.4, window = 100000, step = 13000, readCountThreshold = 20, contaminationAdjustment = “TRUE”, minMappabilityPerWindow = 0.95, breakPointType = 4, minCNAlength = 1) was conducted using known alterations as reference. The concatenation of adjacent CNAs was done using the “merge” option of Bedtools (v2.25.0)^63^. For each gene located in identified CNA regions, a median log2 copy ratio (log2R) was then calculated by considering small windows at least partly overlapping a segment centered on the gene and extended by 25 kb on each side.

### RNA sequencing and analysis

RNA-seq libraries were constructed according to TruSeq Protocols (Illumina) and sequencing was performed using an Illumina HiSeq 2000/4000 instrument. Trimming of sequencing adapters and low-quality bases was done using the Trimmomatic (v0.38) tool^64^. Resulting reads were mapped to the reference using the RNA-seq aligner STAR (v2.7.1)^65^ and quantification of gene and isoform expression were performed using RSEM (v1.3.2)^66^. The limma package and its Voom method^67^ was used to conduct differential expression analysis. Point mutations were identified from RNAseq data as previously described^10^. The GSEA software^68^ was used to conduct the gene set enrichment analysis (GSEA). Genes were pre-ranked according to the signed p-values obtained from the differential expression analysis (sign(logFC)*-log10(P-Value)). The Wikipathways and hallmark sets from the Human Molecular Signatures Database (MSigDB) were used for the analysis.

### Ribosomal eigengene expression calculation

The principal components (PCs) of log2-transformed, normalized expression data (TPM) for RPGs (KEGG gene sets: https://www.gsea-msigdb.org/gsea/msigdb/cards/KEGG_RIBOSOME) were calculated using the prcomp function in R. The ribosomal eigengene was defined as the value of the first principal component (PC1) for each sample. Eigengene expression values were then rescaled to generate a vector ranging from -1 to 1. Note that ribosomal eigengene values did not rely on the expression of RPGs that were found as commonly deleted in the -3/del(3p) group (**Sup. Figure S8**).

### Translational level assay

Puromycin (10 μg/ml) was incorporated in primary specimens for 30 min. Cells were then fixed, permeabilized and intracellular staining of puromycin was achieved using Alexa Fluor 647 labeled anti-puromycin antibody (Millipore, #MABE343). Relative rates of protein synthesis were assessed by flow cytometry by calculating the geometric mean fluorescence intensity of puromycin signals after subtracting background fluorescence for each specimen.

### High-throughput screening (HTS) assay

HTS assay of more than 10,000 structurally diverse compounds – including molecules obtained from a pharmaceutical partner (Bristol Myers Squibb) as well as selected groups of molecules proprietary to IRIC – was conducted in 56 primary AML specimens (**Sup. Figure S12**) and 2 normal control samples (expanded CD34+ cells from cord blood samples). Cell culture and viability assays were conducted as previously described^69^. Briefly, for Compound Correlation Clusters (CCCs) determination, percentage of inhibition data for selective hit compounds were rank-transformed and clustered by minimum spanning tree. Groups of molecules in icicle peaks with *σ* > 0.9 were selected and further filtered for elimination of outlier compounds by selection of profiles correlating with *r* > 0.9 to the median of the group. The remaining compounds were selected as part of CCCs.

### Validation chemical screen

Freshly thawed primary AML specimens were used for chemical screens following the procedure previously described by our group^70^. Compounds were dissolved in DMSO at 10 mM, diluted in media immediately before use and added to seeded cells in serial dilution to determine IC_50_ values (8 dilutions, 1:3, 10 μM down to 4.5 nM) or at the unique concentration of 1 μM, in duplicate wells. Cell viability was evaluated after 6 days in culture using the CellTiterGlo assay (Promega), and compared to DMSO-treated cells.

### Western blots

Alvespimicin (Selleckchem), Geldanamycin (MedChemExpress) and 17-AAG (MedChemExpress) were used at 1M for U937 cells and 500nM for primary specimens. U937 cells were treated with indicated compounds for 24hrs. For western blot analysis, U937 cells were harvested and lysed in total protein extraction lysis buffer (25mM tris ph7.5, 150mM NaCl, 1% NP-40, protease inhibitors). Anti-human antibodies were used to detect RPS14 (ThermoFisher, #PA5-77004), RPL14 (ThermoFisher, #A305-051A), RPL29 (ThermoFisher, #PA5-118248) and Tubulin (TUBA, Cell Signaling Technology, #2144).

### AML specimen treatment

AML cells were thawed at 37 °C in Iscove’s modified Dulbecco’s medium (IMDM) containing 20% FBS and DNase I (100 μg/ml). Cells were cultured for 4hrs in presence or absence of HSP90 inhibitors (Alvespimycin, Geldanamycin and 17-AAG, 500nM) in IMDM, 15% BIT (bovine serum albumin, insulin, transferrin; Stem Cell Technologies 09500), 100 ng/ml SCF (Shenandoah 100-04), 50 ng/ml FLT3L (Shenandoah 100-21), 20 ng/ml Il-3 (Shenandoah 100-80), 20 ng/ml G-CSF (Shenandoah 100-72), 10^−4^ M β-mercaptoethanol, gentamicin (50 μg/ml) and ciprofloxacin (10 μg/ml). Cells were then subjected to Western Blot analysis.

### Statistical analysis

Statistical analyses of all experiments were done using R (version 4.3.2). Depending on the dataset, Fisher’s Exact test, Student’s t-test or Mann-Whitney test were used to determine significance (p-value < 0.05). All p-values lower than 1e-04 were indicated as “p < 1e-04” in the figures (the annotation “p << 1e-04” was used to indicate particularly strong associations).

## Sup. figure legends

**Sup. Figure S1**. General mutation heatmap of the Leucegene cohort. AML-associated genes (y-axis) composing the heatmap were either mutated in one or more specimens (x-axis). Genes were ordered (from top to bottom) based on their mutation frequencies (indicated by the yellow bar graph on the left). Specimens were grouped according to their mutation status (from left to right). SNV: single-nucleotide variant, INS: insertion, ITD: internal tandem duplication, DEL: deletion. See legend at the bottom right for the color code.

**Sup. Figure S2**. Median log2 copy ratio of the common deleted regions (CDR) on chromosomes 3p (3p22.1-3p13) and 5q (5q31.1-5q31.3) (**left**) and variant allele frequencies (VAFs) of homozygous *TP53* mutations in -3/del(3p) specimens (**right**).

**Sup. Figure S3**. Kaplan meier curves for overall survival in patients from the Leucegene AML prognostic cohort (n = 470) with -3/del(3p) (n = 13), -5/del(5q) without 3p alteration (n = 33) or other AML (n = 424). HR: hazard ratio, HSCT: Hematopoietic stem cell transplantation, OS: overall survival.

**Sup. Figure S4**. Pairwise correlation heatmaps of cytosolic RPGs in -3/del(3p) and other AML. Color code (indicated on the right side of each plot) is representative of Pearson’s correlation coefficients (r).

**Sup. Figure S5**. Correlation between *RPL15* gene expression and other cytosolic RPGs in -3/del(3p) specimens (x-axis) or other AMLs (y-axis). Dot colors are representative of the log fold change (logFC) calculated from the differential expression analysis presented in Figure 3A.

**Sup. Figure S6**. Translational levels (assessed by puromycin labeling) of a -3/del(3p) (orange dots) and control AML (blue dots), presenting low and high ribosomal eigengene values respectively. The Pearson’s correlation coefficient is indicated in the figure. The dotted black line results from a least squares regression. MFI, mean fluorescence intensity.

**Sup. Figure S7**. Cytosolic ribosomal eigengene expression values according to *RPL15* (3p24.2) and *RPS14* (5q33.1) expression levels ((-) and (+) symbols depict low (quartile 1 of the whole Leucegene cohort) and high (quartiles 3 and 4) levels of expression, respectively). P-values resulting from the tests comparing each group to the -3/del(3p) AML are directly indicated on the figure (* p-value obtained for the comparison between the test and each control group under the transversal bar).

**Sup. Figure S8**. Comparison of ribosomal eigengene vectors calculated by including (x-axis) or excluding (y-axis) RPGs identified as commonly deleted in -3/del(3p) specimens (i.e. *RPL14, RPL15, RPL29, RPS14, RPSA, RPL7P1, RPL32, RPL26L1, RPS23*).

**Sup. Figure S9**. Frequency of RPG-CNAs per specimen composing each group (as indicated at the top each plot) for each autosome. Autosomes with a null frequency were not reported. Groups are mutually exclusive.

**Sup. Figure S10**. Barplot representation of the percentage of hemizygous *TP53*-mutated -3/del(3p) (orange bars) and -5/del(5q) (blue bars) specimens for 5q cytobands (left axis). The black curve indicates the p-values resulting from Fisher’s exact tests comparing -3/del(3p) to -5/del(5q) AML for each cytoband (right axis). The horizontal dashed line indicates a p-value of 0.05.

**Sup. Figure S11. A**. Correlation between hallmark sets of the Human Molecular Signatures Database (MSigDB) and the ribosomal eigengene expression (Pearson’s correlation). Black dots correspond to outlier genes. The solid blue line corresponds to the median correlation obtained for the 50 hallmark sets together. **B**. Volcano plot representation of enrichments obtained from the gene set enrichment analysis (GSEA) comparing -3/del(3p) AML to other AML and using MYC targets and PI3K-AKT-MTOR pathways from the hallmark sets of the Human Molecular Signatures Database (MSigDB). NES: normalized enrichment score. Blue dots depict significant hits with negative NES. **C**. Ribosomal eigengene expression according to *NPM1* mutation status. Median values are indicated by black lines on each dotplot. NS, non-significant.

**Sup Figure S12**. Cytogenetic subgroup distribution of the 56 AML samples tested in the high-throughput screening assay of 10,000 compounds. CK: complex karyotype, HD: hyperdiploid, IAK: intermediate abnormal karyotype (except isolated trisomy/tetrasomy 8), NK: normal karyotype, T8: tri/tetrasomy 8.

## Acknowledgements

The authors wish to thank Muriel Draoui for Leucegene project coordination and the coinvestigators and members of the Quebec Leukemia Cell Bank. J.F.S. was supported by an IVADO and Canada First Research Excellence Fund (Apogée/CFREF) and a Canadian Institutes of Health Research (CIHR) postdoctoral fellowships. This work was supported by the Government of Canada through Genome Canada and the Ministère de l’économie et de l’innovation du Québec through Génome Québec (ref. grant #4524 and grant #13528). J.H. holds a leukemia research chair from Industrielle-Alliance (Université de Montréal). The Quebec Leukemia Cell Bank is supported by grants from the Cancer Research Network of the Fonds de recherche du Québec–Santé. RNA-Seq read mapping and transcript quantification were performed on the supercomputer Briaree from Université de Montréal, managed by Calcul Québec and Compute Canada. The operation of this supercomputer is funded by the Canada Foundation for Innovation (CFI), NanoQuébec, RMGA and the Fonds de recherche du Québec - Nature et technologies (FRQ-NT).

## Author Contributions

J.F.S. designed the project, analyzed the data, generated all figures (with the exception of the Sup. Figure S1), and wrote the manuscript. J.C. designed experiments testing the sensitivity to HSP90 inhibitors, produced the western blots and, along with I.B., conducted the translational level assay. I.B. conducted the validation chemical screen. G.R.C. conducted the survival analysis. C.M. and N.M. contributed to the high-throughput screening experiment or processing. F.B. was involved in the processing of clinical data of the Leucegene cohort. V.P.L. contributed to Leucegene data processing/acquisition and small mutation analysis. J.H. contributed to project conception, provided AML samples and clinical data of the Leucegene cohort, and analyzed cytogenetic information. G.S. contributed to the project conception and coordination.

## Data Availability

DNA and RNA sequences of the 691 leukemic samples of the Leucegene AML cohort have been deposited in GEO (accession number GSE232130) and SRA (accession number SUB9364031).

Non-identifiable clinical data for the Leucegene AML patient cohort is available to academic investigators with research ethics committee approval in accordance with the procedures of the Quebec Leukemia Cell Bank: [Banque de cellules leucémiques du Québec - Request for Cells/Data](https://bclq.org/request-for-cells/) (bclq.org).

