## Supplemental Figures for "*TP53*-mutant AML with ribosomal gene loss exhibits impaired protein translation and sensitivity to HSP90 inhibition"

### Figure S1

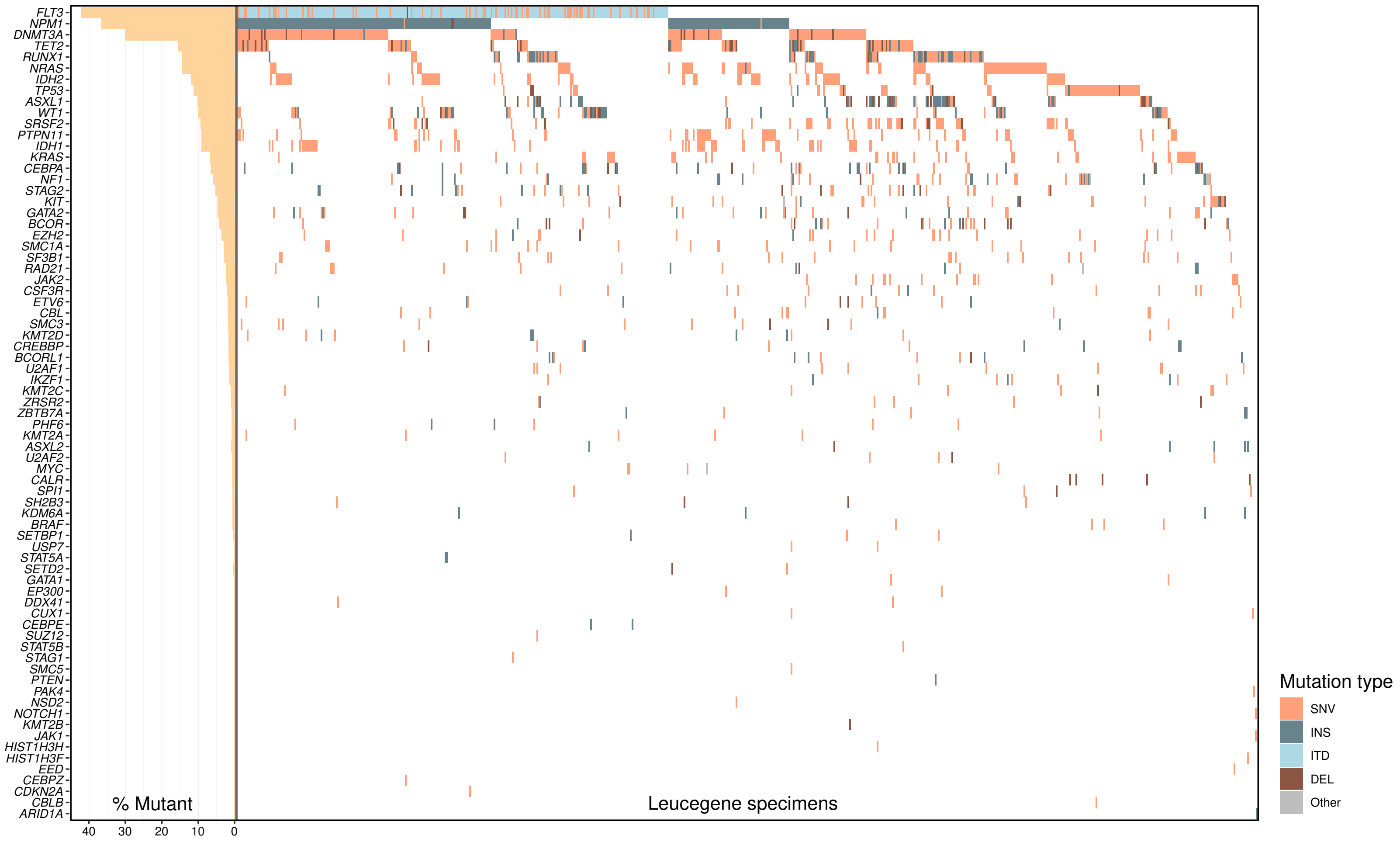
FLT3
NPM1
DNMT3A
TET2
RUNX1
NRAS
IDH2
TP53
ASXL1
WT1
SRSF2
PTPN11
IDH1
KRAS
CEBPA
NF1
STAG2
KIT
GATA2
BCOR
EZH2
SMC1A
SF3B1
RAD21
JAK2
CSF3R
ETV6
CBL
SMC3
KMT2D
CREBBP
BCORL1
U2AF1
IKZF1
KMT2C
ZRSR2
ZBTB7A
PHF6
KMT2A
ASXL2
U2AF2
MYC
CALR
SPI1
SH2B3
KDM6A
BRAF
SETBP1
USP7
STAT5A
SETD2
GATA1
EP300
DDX41
CUX1
CEBPE
SUZ12
STAT5B
STAG1
SMC5
PTEN
PAK4
NSD2
NOTCH1
KMT2B
JAK1
HIST1H3H
HIST1H3F
EED
CEBPZ
CDKN2A
CBLB
ARID1A
% Mutant
40
30
20
10
0
Leucegene specimens
Mutation type
SNV
INS
ITD
DEL
Other

Figure S2

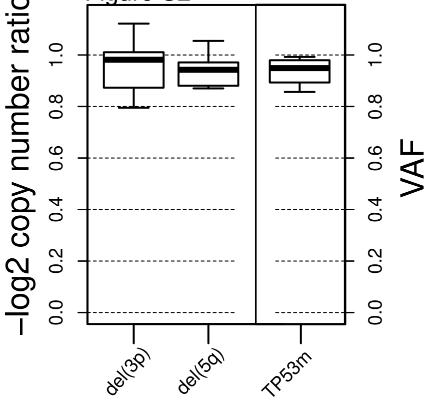

-log2 copy number ratio
0.0
0.2
0.4
0.6
0.8
1.0
VAF
0.0
0.2
0.4
0.6
0.8
1.0
del(3p)
del(5q)
TP53m

Figure S3

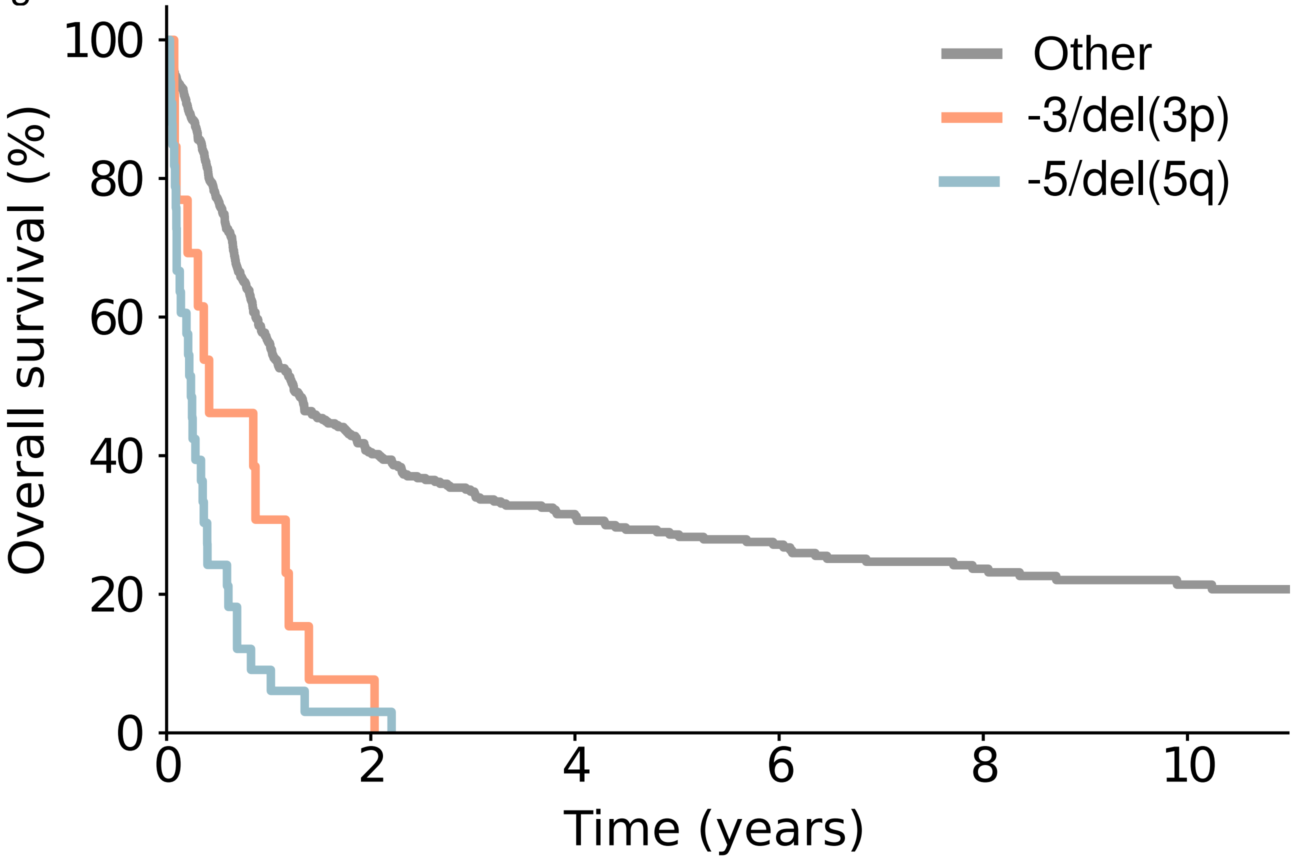


Number at risk

| | | | | | | |
|---|---|---|---|---|---|---|
| Other | 424 | 154 | 99 | 69 | 46 | 32 |
| -3/del(3p) | 13 | 1 | 0 | 0 | 0 | 0 |
| -5/del(5q) | 33 | 1 | 0 | 0 | 0 | 0 |

Figure S4

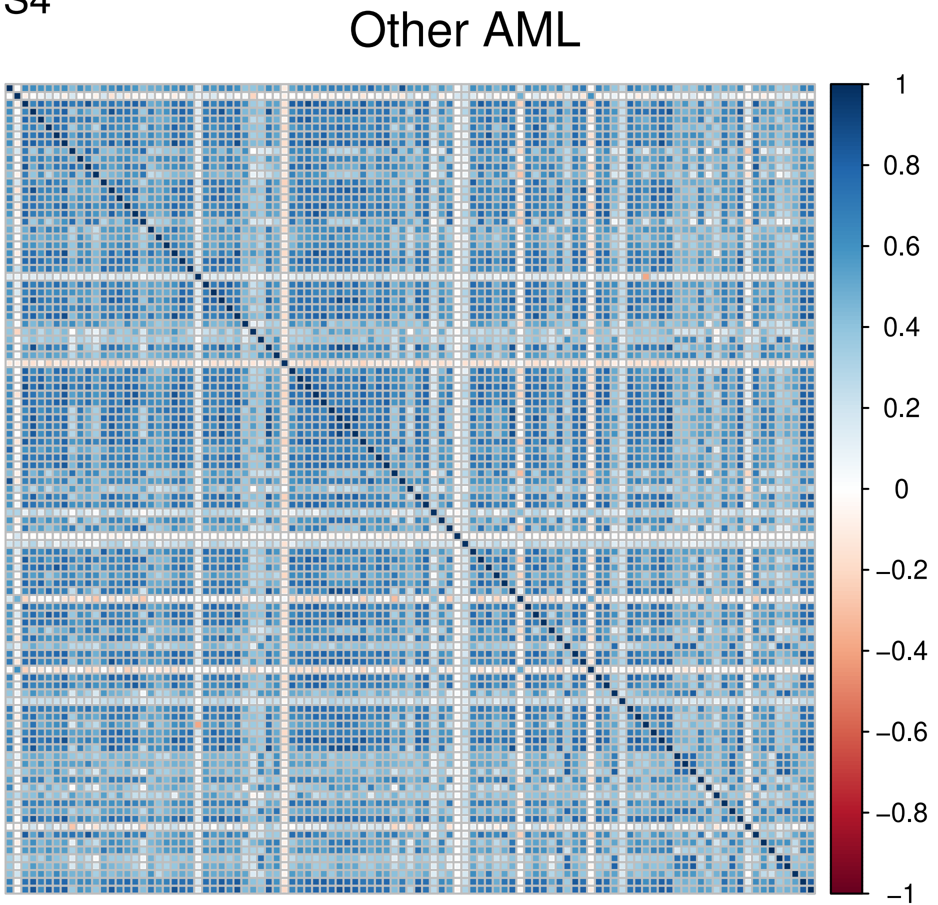

Other AML
1
0.8
0.6
0.4
0.2
0
−0.2
−0.4
−0.6
−0.8
−1


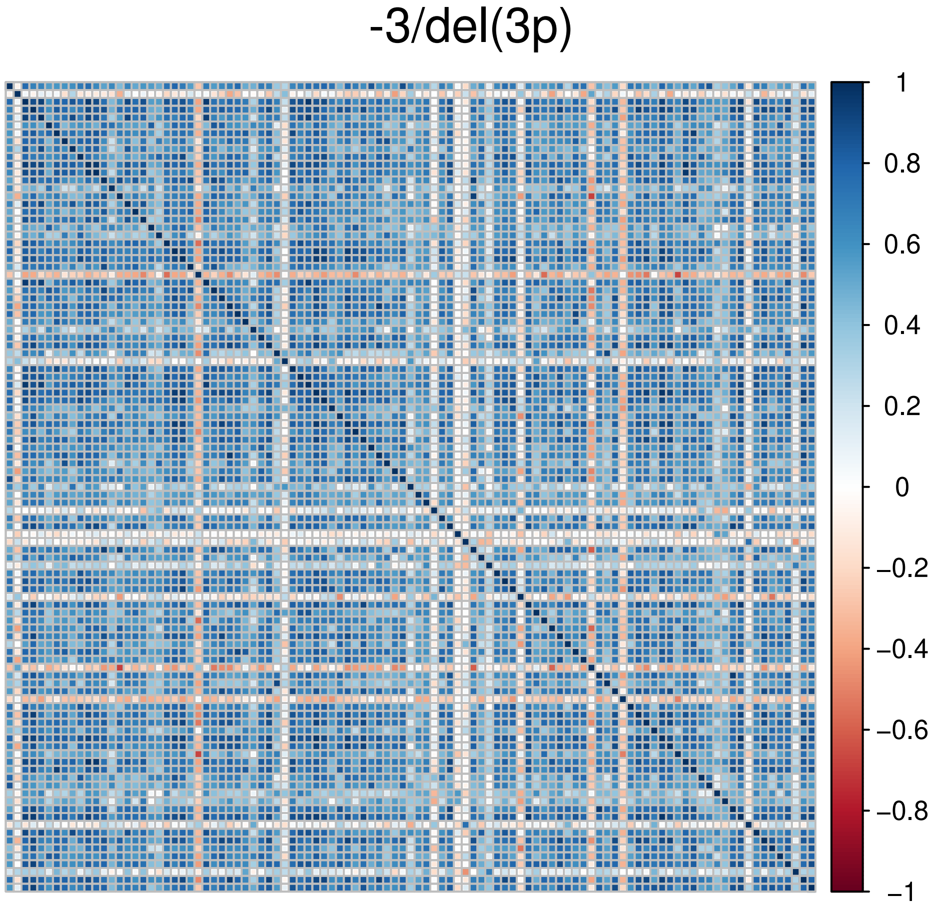

-3/del(3p)
1
0.8
0.6
0.4
0.2
0
−0.2
−0.4
−0.6
−0.8
−1

### Figure S5

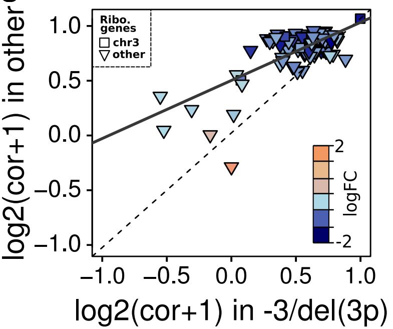

Ribo. genes
chr3
other
logFC
2
-2
log2(cor+1) in other
log2(cor+1) in -3/del(3p)
1.0
0.5
0.0
−0.5
−1.0

Figure S6

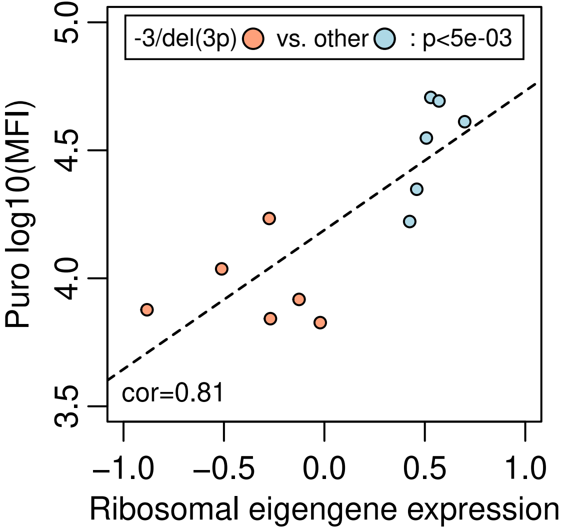
-3/del(3p)
vs. other
: p<5e-03
cor=0.81
Puro log10(MFI)
3.5
4.0
4.5
5.0
−1.0
−0.5
0.0
0.5
1.0
Ribosomal eigengene expression

### Figure S7

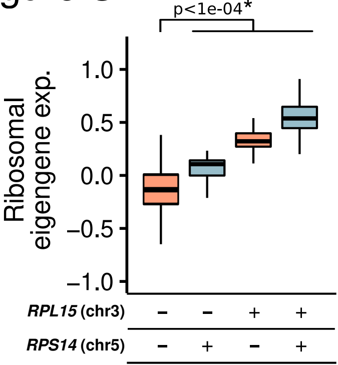
p<1e-04*
Ribosomal eigengene exp.
1.0
0.5
0.0
–0.5
–1.0
RPL15 (chr3)
RPS14 (chr5)
–
–
+
+
–
+
–
+

### Figure S8

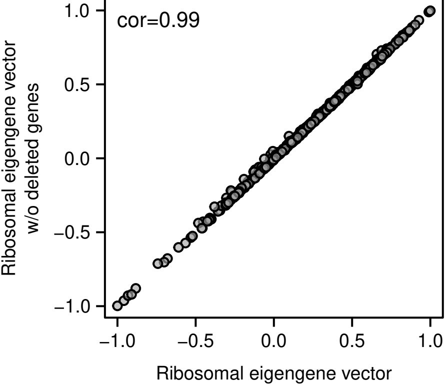
cor=0.99
1.0
0.5
0.0
−0.5
−1.0
Ribosomal eigengene vector
w/o deleted genes
−1.0
−0.5
0.0
0.5
1.0
Ribosomal eigengene vector

Figure S9

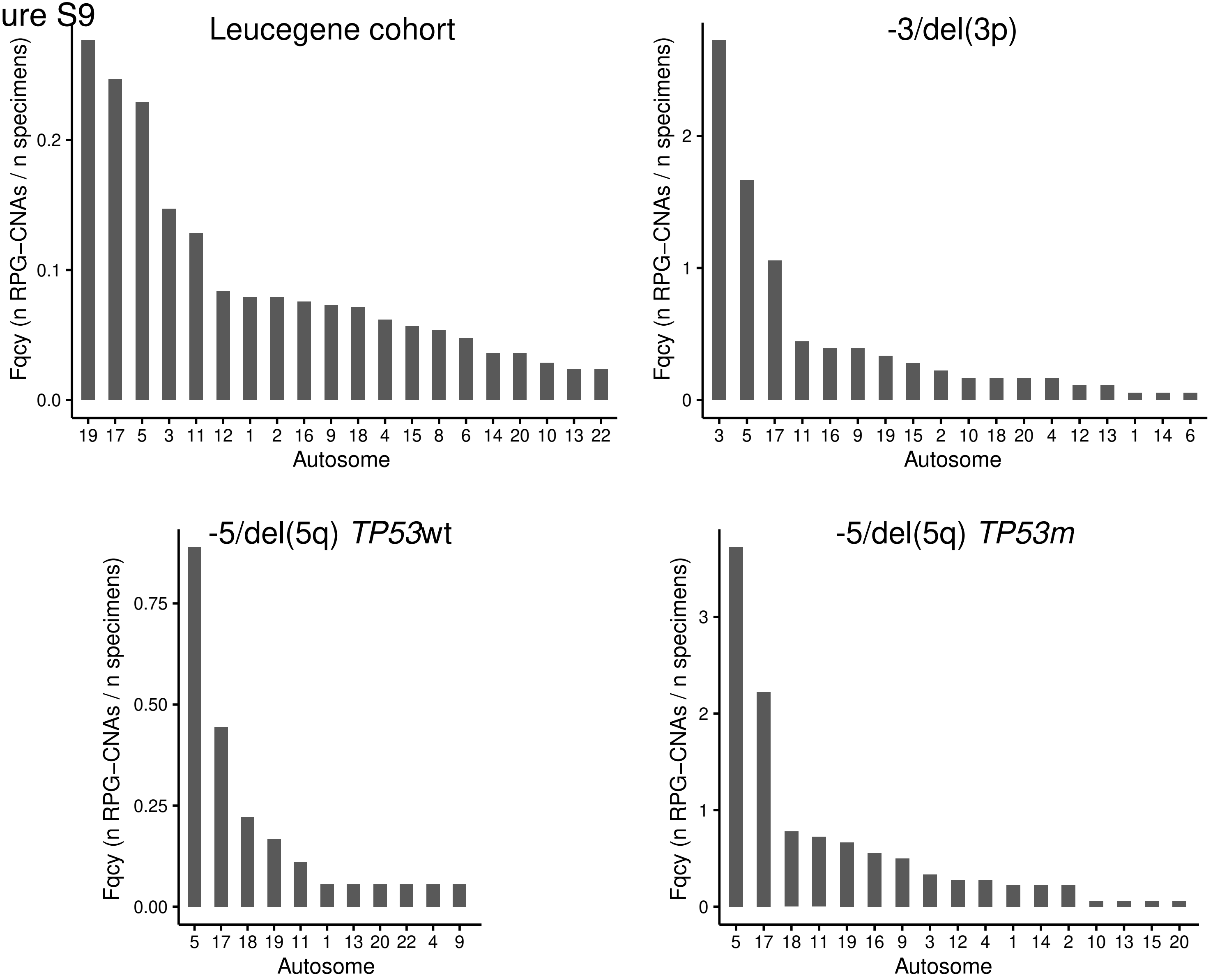
Leucegene cohort
Fqcy (n RPG−CNAs / n specimens)
0.0
0.1
0.2
19 17 5 3 11 12 1 2 16 9 18 4 15 8 6 14 20 10 13 22
Autosome
-3/del(3p)
Fqcy (n RPG−CNAs / n specimens)
0
1
2
3 5 17 11 16 9 19 15 2 10 18 20 4 12 13 1 14 6
Autosome
-5/del(5q) TP53wt
Fqcy (n RPG−CNAs / n specimens)
0.00
0.25
0.50
0.75
5 17 18 19 11 1 13 20 22 4 9
Autosome
-5/del(5q) TP53m
Fqcy (n RPG−CNAs / n specimens)
0
1
2
3
5 17 18 11 19 16 9 3 12 4 1 14 2 10 13 15 20
Autosome

Figure S10

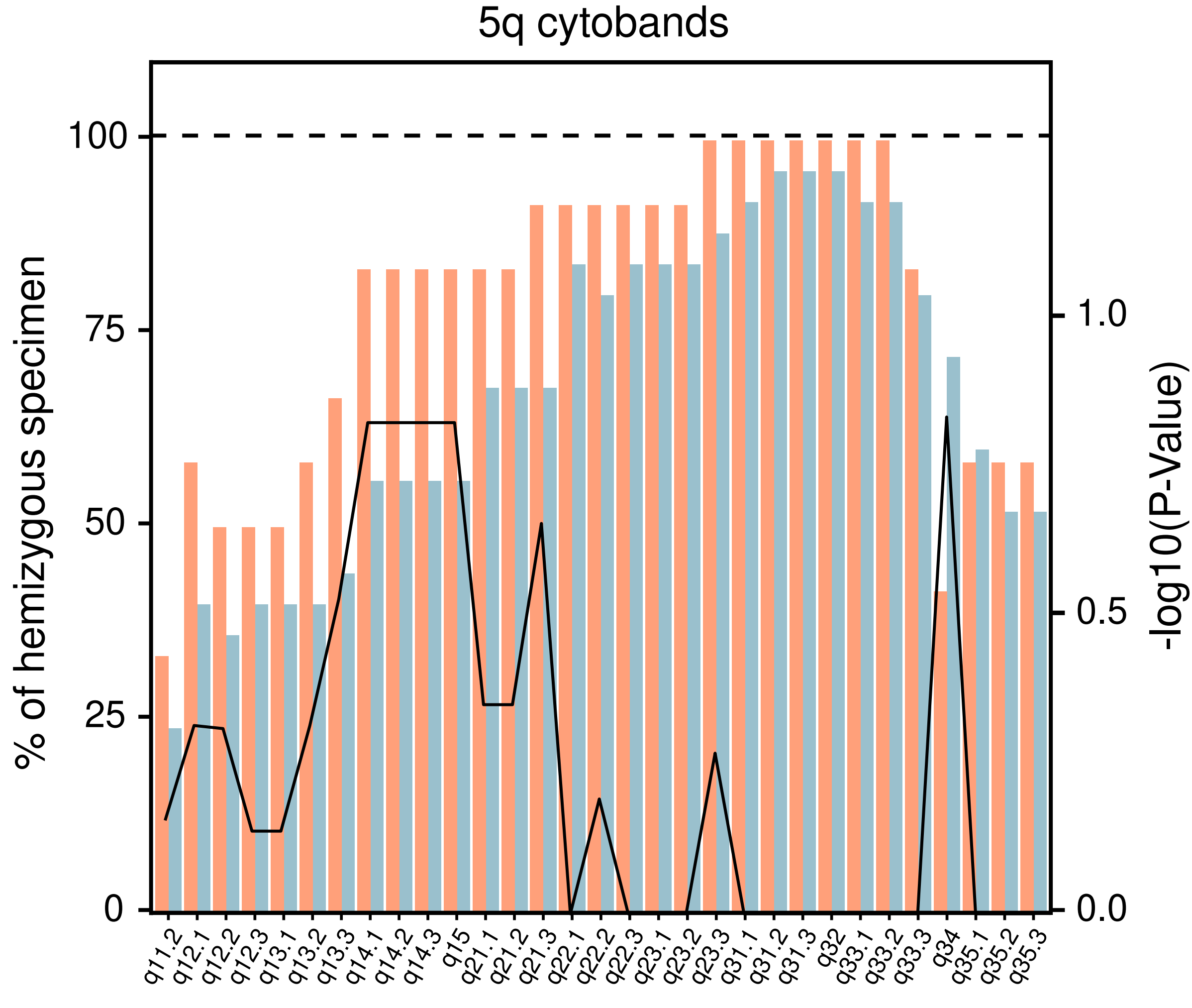
5q cytobands
% of hemizygous specimen
-log10(P-Value)
100
75
50
25
0
1.0
0.5
0.0
q11.2
q12.1
q12.2
q12.3
q13.1
q13.2
q13.3
q14.1
q14.2
q14.3
q15
q21.1
q21.2
q21.3
q22.1
q22.2
q22.3
q23.1
q23.2
q23.3
q31.1
q31.2
q31.3
q32
q33.1
q33.2
q33.3
q34
q35.1
q35.2
q35.3

Figure S11

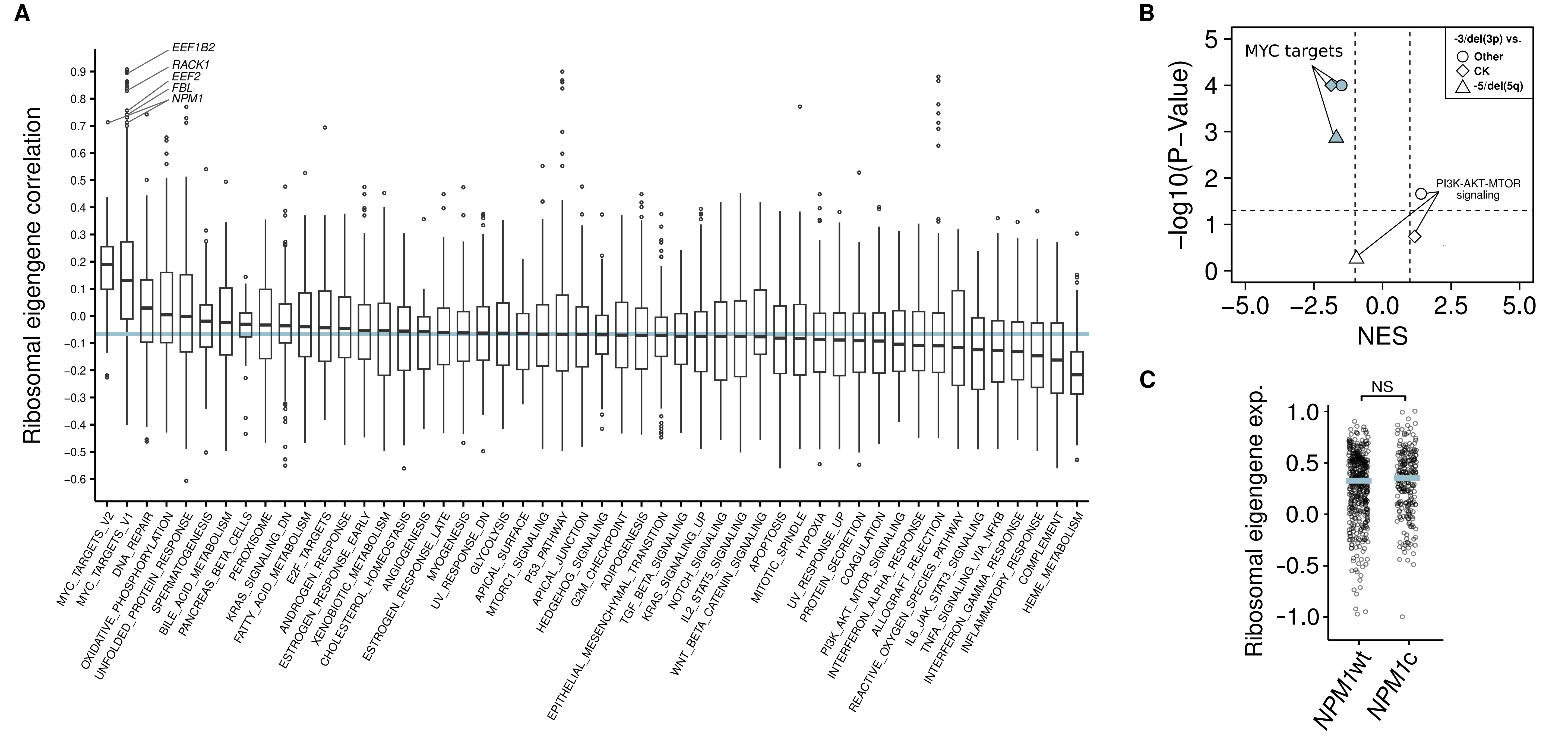

A
Ribosomal eigengene correlation
EEF1B2
RACK1
EEF2
FBL
NPM1
MYC_TARGETS_V2
MYC_TARGETS_V1
DNA_REPAIR
OXIDATIVE_PHOSPHORYLATION
UNFOLDED_PROTEIN_RESPONSE
SPERMATOGENESIS
BILE_ACID_METABOLISM
PANCREAS_BETA_CELLS
PEROXISOME
KRAS_SIGNALING_DN
FATTY_ACID_METABOLISM
E2F_TARGETS
ANDROGEN_RESPONSE
ESTROGEN_RESPONSE_EARLY
XENOBIOTIC_METABOLISM
CHOLESTEROL_HOMEOSTASIS
ANGIOGENESIS
ESTROGEN_RESPONSE_LATE
MYOGENESIS
UV_RESPONSE_DN
GLYCOLYSIS
APICAL_SURFACE
MTORC1_SIGNALING
P53_PATHWAY
APICAL_JUNCTION
HEDGEHOG_SIGNALING
G2M_CHECKPOINT
ADIPOGENESIS
EPITHELIAL_MESENCHYMAL_TRANSITION
TGF_BETA_SIGNALING
KRAS_SIGNALING_UP
NOTCH_SIGNALING
IL2_STAT5_SIGNALING
WNT_BETA_CATENIN_SIGNALING
APOPTOSIS
MITOTIC_SPINDLE
HYPOXIA
UV_RESPONSE_UP
PROTEIN_SECRETION
COAGULATION
PI3K_AKT_MTOR_SIGNALING
INTERFERON_ALPHA_RESPONSE
ALLOGRAFT_REJECTION
REACTIVE_OXYGEN_SPECIES_PATHWAY
IL6_JAK_STAT3_SIGNALING
TNFA_SIGNALING_VIA_NFKB
INTERFERON_GAMMA_RESPONSE
INFLAMMATORY_RESPONSE
COMPLEMENT
HEME_METABOLISM
B
-log10(P-Value)
NES
MYC targets
PI3K-AKT-MTOR signaling
-3/del(3p) vs.
Other
CK
-5/del(5q)
C
Ribosomal eigengene exp.
NS
NPM1wt
NPM1c

### Figure S12

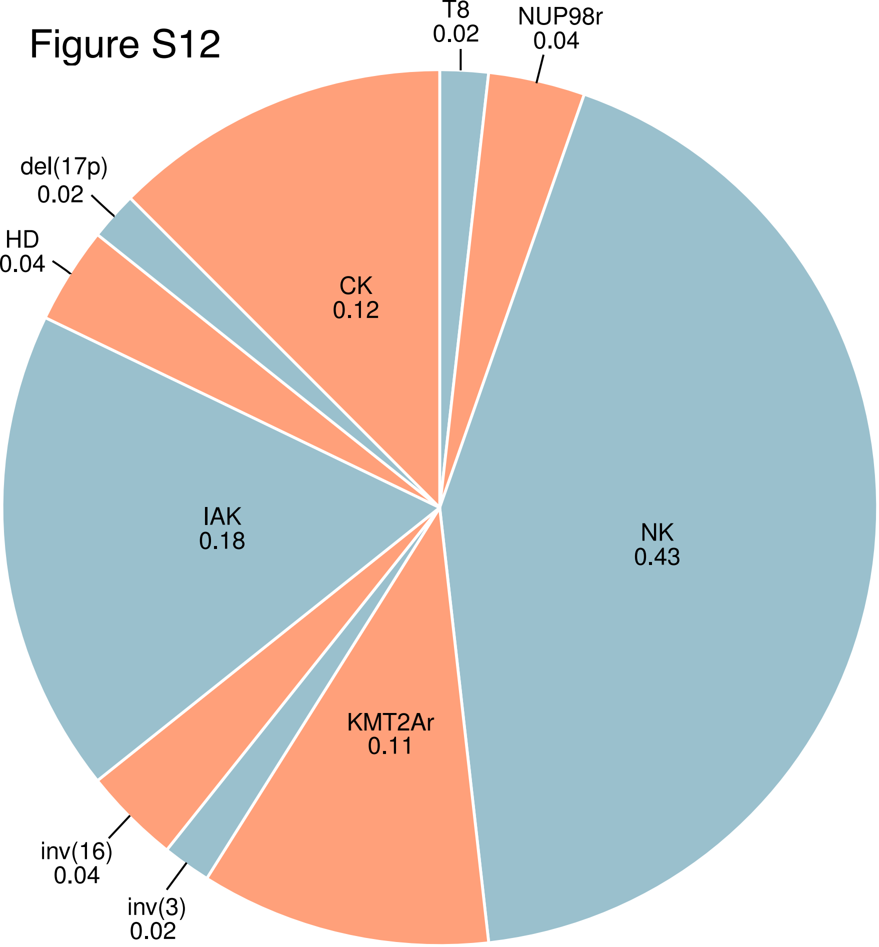
T8
0.02
NUP98r
0.04
NK
0.43
KMT2Ar
0.11
inv(3)
0.02
inv(16)
0.04
IAK
0.18
HD
0.04
del(17p)
0.02
CK
0.12
